# Optimal anticipatory control as a theory of motor preparation: a thalamo-cortical circuit model

**DOI:** 10.1101/2020.02.02.931246

**Authors:** Ta-Chu Kao, Mahdieh S. Sadabadi, Guillaume Hennequin

## Abstract

Across a range of motor and cognitive tasks, cortical activity can be accurately described by low-dimensional dynamics unfolding from specific initial conditions on every trial. These “preparatory states” largely determine the subsequent evolution of both neural activity and behaviour, and their importance raises questions regarding how they are — or ought to be — set. Here, we formulate motor preparation as optimal anticipatory control of future movements, and show that the solution requires a form of internal feedback control of cortical circuit dynamics. In contrast to a simple feedforward strategy, feedback control enables fast movement preparation and orthogonality between preparatory and movement activity, a distinctive feature of peri-movement activity in reaching monkeys. We propose a circuit model in which optimal preparatory control is implemented as a thalamo-cortical loop gated by the basal ganglia.

---

Fast ballistic movements (e.g. throwing) require spatially and temporally precise commands to the musculature. Many of these signals are thought to arise from internal dynamics in the primary motor cortex (M1; Figure 1A; Evarts, 1968; Todorov, 2000; Scott, 2012; Shenoy et al., 2013; Omrani et al., 2017). In turn, consistent with state trajectories produced by a dynamical system, M1 activity during movement depends strongly on the “initial condition” reached just before movement onset, and variability in initial condition predicts behavioural variability (Churchland et al., 2006; Afshar et al., 2011; Pandarinath et al., 2018). An immediate consequence of this dynamical systems view is the so-called “optimal subspace hypothesis” (Churchland et al., 2010; Shenoy et al., 2013): the network dynamics that generate movement must be seeded with an appropriate initial condition prior to each movement. In other words, accurate movement production likely requires fine adjustment of M1 activity during a phase of movement preparation (Figure 1B, green).

**Figure 1:**
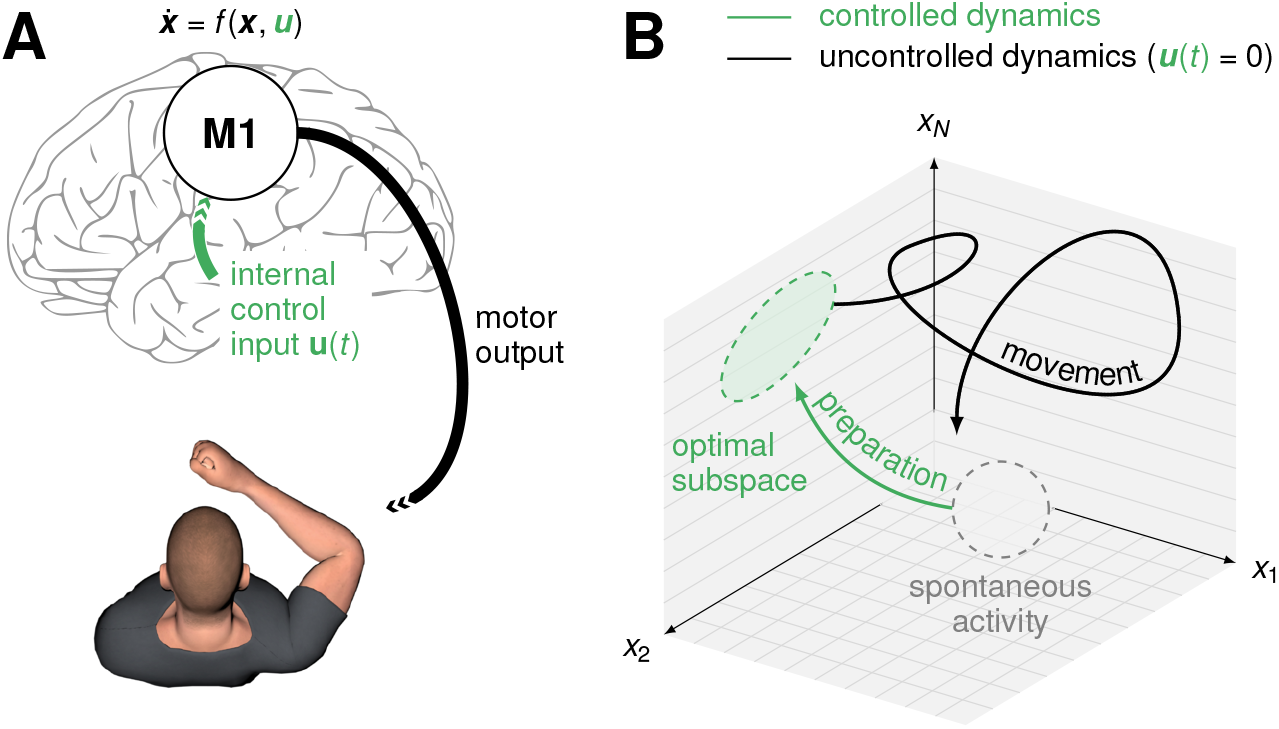
Preparation & execution of ballistic movements. (**A**) Under a dynamical systems view of motor control (Shenoy et al., 2013), movement is generated by M1 dynamics. Prior to movement, the M1 population activity state ***x***(*t*) must be controlled into an optimal, movement-specific subspace in a phase of preparation; this requires internally generated control inputs ***u***(*t*). (**B**) Schematic state space trajectory during movement preparation and execution.

The optimal subspace hypothesis helps to make sense of neural activity during the preparation epoch, yet several unknowns remain. What should the structure of the optimal preparatory subspace be? How does this structure depend on the dynamics of the cortical network during the movement epoch, and on downstream motor processes? Must preparatory activity converge to a single movement-specific state and be held there until movement initiation, or is some slack allowed? What are the dynamical processes and associated circuit mechanisms responsible for motor preparation? These questions can be (and have been partially) addressed empirically, e.g. through analyses of neural population recordings in reaching monkeys (Churchland et al., 2010; Ames et al., 2014; Elsayed et al., 2016) or optogenetic dissection of circuits involved in motor preparation (Li et al., 2016; Guo et al., 2017; Gao et al., 2018; Sauerbrei et al., 2020). Yet, for lack of an appropriate theoretical scaffold, it has been difficult to interpret these experimental results within the broader computational context of motor control.

Here, we bridge this gap by considering motor preparation as an integral part of motor control. We show that optimal control theory, which has successfully explained behaviour (Todorov and Jordan, 2002; Scott et al., 2015) and neural activity (Todorov, 2000; Lillicrap and Scott, 2013) during the movement epoch, can also be brought to bear on motor preparation. Specifically, we argue that there is a prospective component of motor control that can be performed in anticipation of the movement (i.e. during preparation). This leads to a normative formulation of the optimal subspace hypothesis. Our theory specifies the control inputs that must be given to the movement-generating network during preparation to ensure that (i) any subsequent motor errors are kept minimal and (ii) movements can be initiated rapidly. These optimal inputs can be realized by a feedback loop onto the cortical network.

This normative model provides a core insight: the “optimal subspace” is likely high dimensional, with many different initial conditions giving rise to the same correct movement. This has an important consequence for preparatory control: at the population level, only a few components of preparatory activity impact future motor outputs, and it is these components only that need active controlling. By taking this into account, the optimal preparatory feedback loop dramatically improves upon a simpler feedforward strategy. This holds for multiple classes of network models trained to perform reaches, and whose movement-epoch dynamics are quantitatively similar those of monkey M1. Moreover, optimal feedback inputs, but not feedforward inputs, robustly orthogonalize preparatory- and movement-epoch activity, thus accounting for one of the most prominent features of perimovement activity in reaching monkeys (Kaufman et al., 2014; Elsayed et al., 2016).

Finally, we propose a way in which neural circuits may implement optimal anticipatory control. In particular, we propose that cortex is actively controlled by thalamic feedback during motor preparation, with thalamic afferents providing the desired optimal control inputs. This is consistent with the causal role of thalamus in the preparation of directed licking in mice (Guo et al., 2017). Moreover, we posit that the basal ganglia operate an on/off switch on the thalamocortical loop (Jin and Costa, 2010; Cui et al., 2013; Halassa and Acsády, 2016; Logiaco et al., 2019), thereby flexibly controlling the timing of both movement planning and initiation.

Beyond motor control, a broader set of cortical computations are also thought to rest on low dimensional circuit dynamics, with initial conditions largely determining behaviour (Pandarinath et al., 2018; Sohn et al., 2019). These computations, too, may hinge on careful preparation of the state of cortex in appropriate subspaces. Our framework, and control theory more generally, may provide a useful language for reasoning about putative algorithms and neural mechanisms (Kao and Hennequin, 2019).

## Results

### A model of movement generation

We begin with an inhibition-stabilized network (ISN) model of the motor cortex in which a detailed balance of excitation and inhibition enables the production of rich, naturalistic activity transients (STAR Methods; Hennequin et al., 2014, Figure 2A). This network serves as a pattern generator for the production of movement (we later investigate other movement-generating networks). Specifically, the network directly controls the two joint torques of a two-link arm, via a linear readout of the momentary network firing rates:

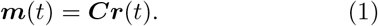

**Figure 2:**
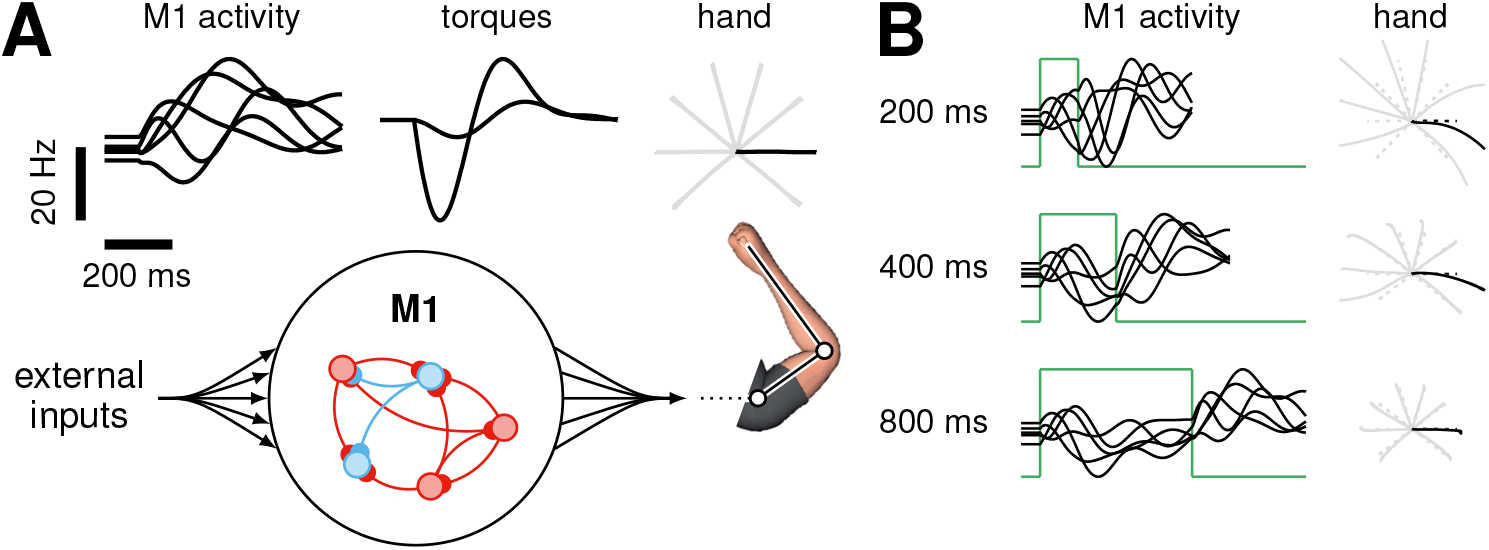
Movement generation in a network model of M1. **(A)** Schematics of our M1 model of motor pattern generation. The dynamics of an excitation-inhibition network (Hennequin et al., 2014) unfold from movement-specific initial conditions, resulting in firing rate trajectories (left; 5 neurons shown) which are linearly read out into joint torques (middle), thereby producing hand movements (right). The model is calibrated for the production of eight straight center-out reaches (20 cm length); firing rates and torques are shown only for the movement colored in black. To help visualize initial conditions, firing rates are artificially clamped for the first 100 ms. **(B)** Network activity and corresponding hand trajectories as in A, for three different preparation lengths, under the naive feedforward strategy whereby a static input step (green) moves the fixed point of the dynamics to the desired initial condition.

Here, ***m***(*t*) is a vector containing the momentary torques, and ***r***(*t*) is the population firing rate vector (described below). The network has *N* = 200 neurons, whose momentary internal activations ***x***(*t*) = (*x*_1_, *x*_2_,…, *x_N_*)^*T*^ evolve according to (Dayan and Abbott, 2001):

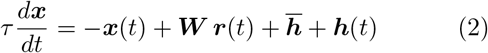

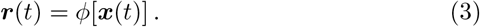

Here, *τ* is the single-neuron time constant, ***W*** is the synaptic connectivity matrix, and *ϕ*[*x*] (applied to ***x*** element-wise) is a rectified-linear activation function converting internal activations into momentary firing rates. The network is driven by two different inputs shared across all movements: a constant input 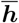 responsible for maintaining a heterogeneous spontaneous activity state ***x***_sp_, and a transient input ***h***(*t*) arising at movement onset and decaying through movement. The latter input models the dominant, conditionindependent timing-related component of monkey M1 activity during movement (Kaufman et al., 2016). We note that, while the network model is generally nonlinear, it can be well approximated by a linear model (***r*** = ***x***) as only a small fraction of neurons are silent at any given time (Figure S5; Discussion). Our formal analyses here rely on linear approximations, but all simulations are based on Equations 2 and 3 with nonlinear *ϕ*.

We calibrated the model for the production of eight rapid straight reaches with bell-shaped velocity profiles (Figure S1). To perform this calibration, we noted that – in line with the dynamical systems view of movement generation (Shenoy et al., 2013) – movements produced by our model depend strongly on the “initial condition”, i.e. the cortical state ***x*** just before movement onset (Churchland et al., 2010; Afshar et al., 2011). We thus “inverted” the model numerically, by optimizing eight different initial conditions and a common readout matrix ***C*** such that the dynamics of the nonlinear model (Equations 2 and 3), seeded with each initial condition, would produce the desired movement. Importantly, we constrained ***C*** so that its nullspace contained the network’s spontaneous activity state, as well as all eight initial conditions. This constraint ensures that movement does not occur spontaneously or during late preparatory stages when ***x***(*t*) has converged to one of these initial conditions. Nevertheless, these constraints are not sufficient to completely silence the network’s torque readout ***m***(*t*) during preparation. While such spurious output tended to be very small in our simulations (Figure S7), the two stages of integration of ***m***(*t*) by the arm’s mechanics led to substantial drift of the hand before the reach. To prevent drift without modelling spinal reflexes for posture control, we artificially set ***m***(*t*) = 0 during movement preparation.

As we show below, our main results do not depend on the details of the model chosen to describe movement-generating M1 dynamics. Here, we chose the ISN model for its ability to produces activity similar to M1’s during reaching, both in single neurons (Figure 2A, top left) and at the population level as shown by jPCA and canonical correlations analysis (Figure S2; Churchland et al., 2012; Sussillo et al., 2015).

### Optimal control as a theory of motor preparation

Having calibrated our network model of movement generation, we now turn to preparatory dynamics. Shenoy et al.’s dynamical systems perspective suggests that accurate movement execution likely requires careful seeding of the generator’s dynamics with an appropriate, reach-specific initial condition (Afshar et al., 2011). In our model, this means that the activity state ***x***(*t*) of the cortical network must be steered towards the initial condition corresponding to the intended movement (Figure 1B, green). We assume that this process is achieved through additional movement-specific control inputs ***u***(*t*) (Figure 1, green):

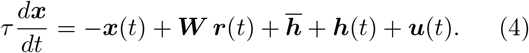

The control inputs ***u***(*t*) are then rapidly switched off to initiate movement.

A very simple way of achieving a desired initial condition ***x***⋆, which we call the “naive feedforward strategy”, is to use a static external input ***u***(*t*) = ***u***⋆ of the form

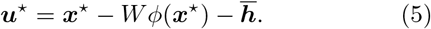

This establishes ***x***⋆ as a fixed point of the population dynamics, as *d**x***/*dt* = 0 in Equation 4 when ***x*** = ***x***⋆. In stable linear networks (and, empirically, in our nonlinear model too), these simple preparatory dynamics are guaranteed to achieve the desired initial state *eventually*, i.e. after sufficiently long preparation time. However, the network takes time to settle in the desired state (Figure 2B); in fact, in a linear network, the response to an input step (here, the feedforward input driving preparation) contains exactly the same timescales as the corresponding impulse reponse (here, the autonomous movement-generating response to the initial condition ***x***⋆). Thus, under the naive feedforward strategy, the duration of the movement itself sets a fundamental limit on how fast ***x***(*t*) can approach ***x***⋆. This is at odds with experimental reports of relatively fast preparation, on the order of ≈ 50 ms, compared to the > 500 ms movement-epoch window over which neural activity is significantly modulated (Lara et al., 2018).

How can preparation be sped up? An important first step towards formalizing preparatory control and unravelling putative circuit mechanisms is to understand how deviations from “the right initial condition” impact the subsequent movement. Are some deviations worse than others? Mathematical analysis reveals that, depending on the direction in state space along which the deviation occurs, there may be strong motor consequences or none at all (Figure 3; STAR Methods). Some preparatory deviations are “prospectively potent”: they propagate through the dynamics of the generator network during the movement epoch, modifying its activity trajectories, and eventually leading to errors in torques and hand motion (Figure 3A, left). Other preparatory deviations are “prospectively readout-null”: they cause subsequent perturbations in cortical state trajectories, too, but these are correlated across neurons in such a way that they cancel in the readout and leave the movement unaltered (Figure 3A, center). Yet other preparatory perturbations are “prospectively dynamic-null”: they are outright rejected by the recurrent dynamics of the network, thus causing little impact on subsequent neuronal activity during movement, let alone on torques and hand motion (Figure 3A, right; Figure 3B-C).

**Figure 3:**
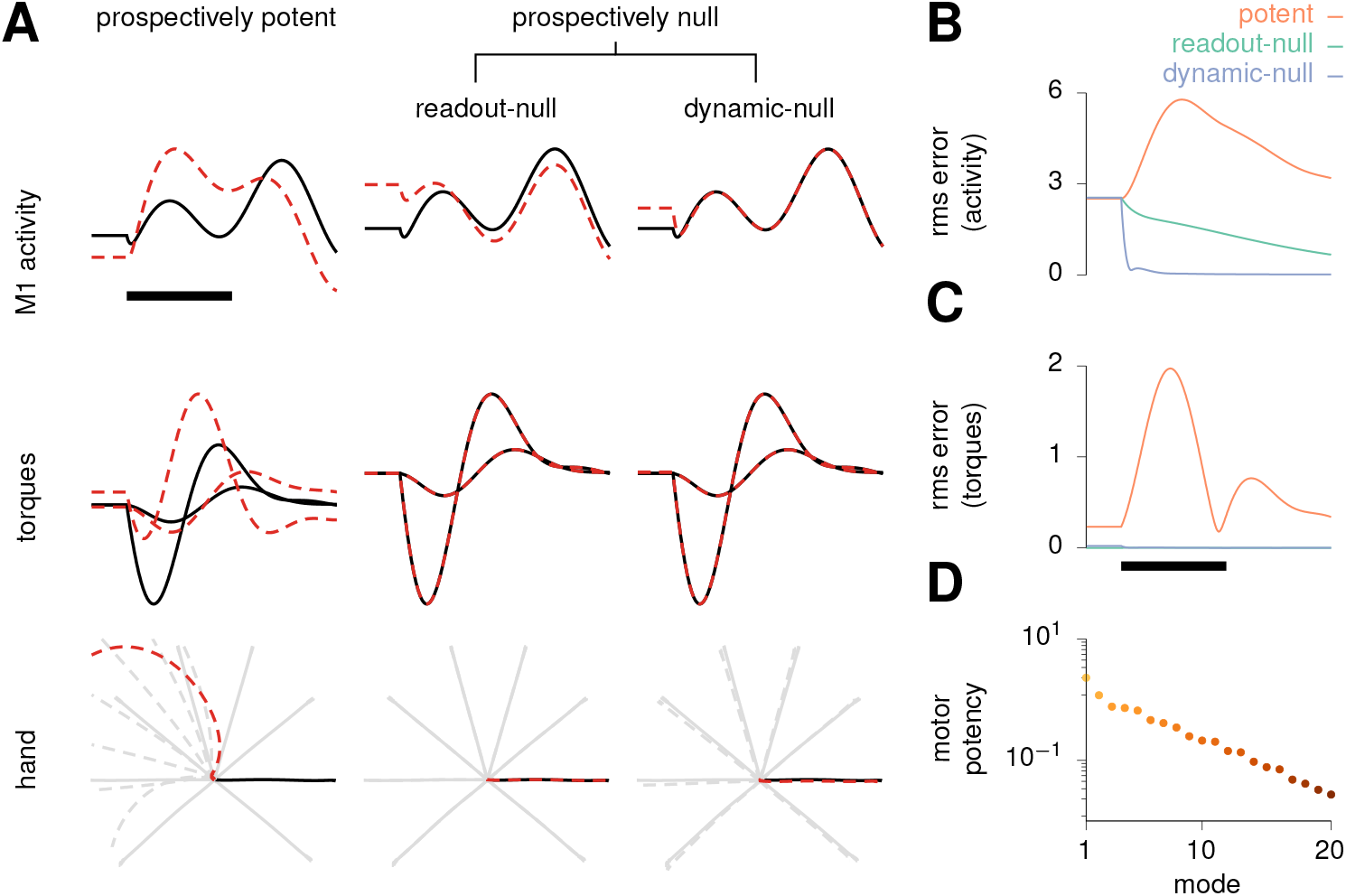
Formalization of the optimal subspace hypothesis. (**A**) Effect of three qualitatively different types of small perturbations of the initial condition (prospectively potent, prospectively readout-null, prospectively dynamic-null) on the three processing stages leading to movement (M1 activity, joint torques, and hand position), as already shown in Figure 2A. Unperturbed traces are shown as solid lines, perturbed ones as dashed red lines. Only one example neuron (top) is shown for clarity. Despite all having the same size here (Euclidean norm), these three types of perturbation on the initial state have very different consequences. Left: “prospectively potent” perturbations result in errors at every stage. Middle: “prospectively readout-null” perturbations cause sizeable changes in internal network activity but not in the torques. Right: “prospectively dynamic-null” perturbations are inconsequential at all stages. **(B)** Timecourse of root-mean-square error in M1 activity across neurons and reach conditions, for the three different types of perturbations. **(C)** Same as (B) but root-mean-square error in torques. **(D)** The motor potency of the top 20 most potent modes. In (A-C), signals are artificially held constant in the first 100 ms for visualization, and black scale bars denote 200 ms from movement onset.

We emphasize that the “prospective potency” of a preparatory deviation is distinct from its “immediate potency”, i.e. from the direct effect such neural activity might have on the output torques (Kaufman et al., 2014; Vyas et al., 2020). For example, the initial state for each movement is prospectively potent by construction, as it seeds the production of movement-generating network responses. However, it does not itself elicit movement on the instant, and so is immediately null. Having clarified this distinction, we will refer to prospectively potent/null directions simply as potent/null directions for succinctness.

The existence of readout-null and dynamic-null directions implies that, in fact, there is no such thing as “the right initial condition” for each movement. Rather, a multitude of initial conditions which differ along null directions give rise to the correct movement. To quantify the extent of this degeneracy, we measure the “prospective potency” of a direction by the integrated squared error in output torques induced during movement by a fixed-sized perturbation of the initial condition along that direction. We can then calculate a full basis of orthogonal directions ranking from most to least potent (an analytical solution exists for linear systems; STAR Methods; Kao and Hennequin, 2019). For our model with only two readout torques, the effective dimensionality of the potent subspace is approximately 8 (Figure 3D; STAR Methods). This degeneracy substantially lightens the computational burden of preparatory ballistic control: there are only a few potent directions in state space along which cortical activity needs active controlling prior to movement initiation. Thus, taking into account the energetic cost of neural control, preparatory dynamics should aim at preferentially eliminating errors in preparatory states along those few directions that matter for movement.

We now formalize these insights in a normative model of preparatory motor control. At any time *t* during preparation, we can assign a “prospective motor error” 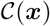 to the current cortical state ***x***(*t*). This prospective error is the total error in movement that *would* result if movement was initiated at this time, i.e. if control inputs were suddenly switched off and the generator network was left to evolve dynamically from ***x***(*t*) (STAR Methods). Note that 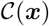 is directly related to the measure of prospective potency described above. An ideal controller would supply the cortical network with such control inputs ***u***(*t*) as necessary to lower the prospective motor error as fast as possible. This would enable accurate movement production in short order. We therefore propose the following cost functional:

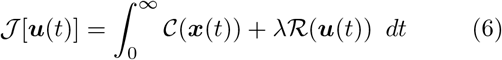

where 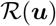 is an energetic cost which penalizes large control signals in excess of a baseline required to hold ***x***(*t*) in the optimal subspace, and λ sets the relative importance of this energetic cost. Note that ***x***(*t*) depends on ***u***(*t*) via Equation 4.

### Optimal preparatory control

When (i) the prospective motor error 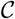 is quadratic in the output torques ***m***, (ii) the energy cost 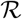 is quadratic in ***u***, and (iii) the network dynamics are linear, then minimizing Equation 6 corresponds to the well-known linear quadratic regulator (LQR) problem in control theory (Skogestad and Postlethwaite, 2007). The optimal solution is a combination of a constant input and instantaneous (linear) state feedback,

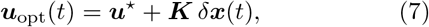

where *δ**x***(*t*) is the momentary deviation of ***x*** from a valid initial condition for the desired movement (STAR Methods). In Equation 7, the constant input ***u***⋆ is movement specific, but the optimal gain matrix ***K*** is generic; both can be derived in algebraic form. Thus, even though the actual movement occurs in “open loop” (without corrective sensory feedback), optimal movement preparation occurs in closed loop, with the state of the pattern generator being controlled via internal feedback in anticipation of the movement. Later in this Results section, we build a realistic circuit model in which optimal preparatory feedback (Equation 7) is implemented as a realistic thalamo-cortical loop. Prior to that, we focus on the core predictions of the optimal control law as an algorithm, independent of its implementation.

Optimal control inputs (Equation 7) lead to multiphasic preparatory dynamics in the cortical network (Figure 4A, top). Single-neuron responses separate across movement conditions in much the same way as they do in the monkey M1 data (Figure 4C), with a transient increase in total across-movement variance (Figure 4D). The prospective motor error decreases very quickly to negligible values (Figure 4A, bottom; note the small green area under the curve) as ***x***(*t*) is driven into the appropriate subspace. After the preparatory feedback loop is switched off and movement begins, the system accurately produces the desired torques and hand trajectories (Figure 4A, middle). Indeed, movements are ready to be performed after as little as 50 ms of preparation (Figure 4B). We note, though, that it is possible to achieve arbitrarily fast preparation by decreasing the energy penalty factor λ in Equation 6 (Figure 4E). However, this is at the price of large energetic costs 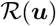, i.e. unrealistically large control inputs.

**Figure 4:**
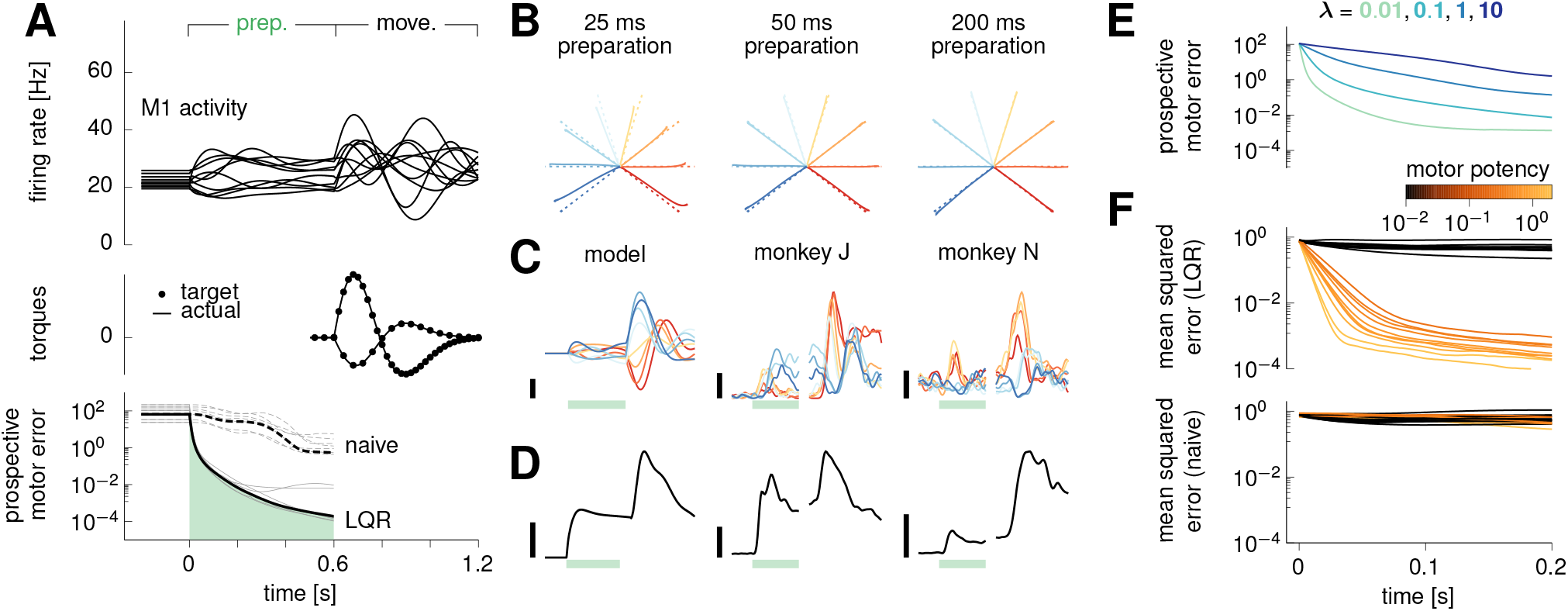
Optimal preparatory control. **(A)** Dynamics of the model during optimal preparation and execution of a straight reach at a 144-degree angle. Optimal control inputs are fed to the cortical network during preparation, and subsequently withdrawn to elicit movement. Top: firing rates of a selection of ten model neurons. Middle: generated torques (line), compared to targets (dots). Bottom: the prospective motor error 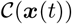 quantifies the accuracy of the movement if it were initiated at time *t* during the preparatory phase. Under the action of optimal control inputs, 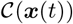 decreases very fast, until it becomes small enough that an accurate movement can be triggered. The dashed line shows the evolution of the prospective cost for the naive feedforward strategy (see text). Gray lines denote the other 7 reaches for completeness. **(B)** Hand trajectories for each of the eight reaches (solid), following optimal preparation over a window of 25 ms (left), 50 ms (center) and 200 ms (right). Dashed lines show the target movements. **(C)** Firing rate of a representative neuron in the model (left), and the two monkeys (center and right) for each movement condition (color-coded as in Figure 2B). Green bars mark the 600 ms preparation window, black scale bars indicate 20 Hz. **(D)** Evolution of the average across-movement variance in single-neuron preparatory activity in the model (left) and the monkeys (center and right). Black scale bars indicate 16 Hz^2^. **(E)** Prospective motor error during preparation, averaged over the eight reaches, for different values of the energy penalty parameter λ. **(F)** The state of the cortical network is artificially set to deviate randomly from the target movement-specific initial state at time *t* = 0, just prior to movement preparation. The temporal evolution of the squared Euclidean deviation from target (averaged over trials and movements) is decomposed into contributions from the 10 most and 10 least potent directions, color-coded by their motor potency as in Figure 3D. In (A-D) and (F), we used λ = 0.1.

The neural trajectories under optimal preparatory control display a striking property, also observed in monkey M1 and PMd recordings (Ames et al., 2014; Lara et al., 2018): by the time movement is ready to be triggered (approx. 50 ms in the model), the firing rates of most neurons have not yet converged to the values they would attain after a longer preparation time — that is, ║*δ**x***(*t*)║ ≫ 0 (Figure 4A and C). Intuitively, this arises for the following reasons. First, the network reacts to the sudden onset of the preparatory input by following the flow of its internal dynamics, thus generating transient activity fluctuations. Second, the optimal controller is not required to suppress the null components of these fluctuations, which have negligible motor consequences. Instead, control inputs are used sparingly to steer the dynamics along potent directions only. Thus, the network becomes ready for movement initiation well before its activity has settled. To confirm this intuition, we artificially set ***x***(*t*) at preparation onset to randomly and isotropically deviate from the target initial condition. We then computed the expected momentary squared deviation *δ**x***(*t*) along the spectrum of potent directions (shown in Figure 3D) during preparation. Errors are indeed selectively eliminated along directions with motor consequences, while they linger or even grow in other, inconsequential directions (Figure 4F, top).

Importantly, feedback control vastly outperforms the naive feedforward strategy of Figure 2B, which corresponds to the limit of zero input energy as it is defined in our framework (STAR Methods; it is also the optimal solution in the limit of λ → ∞ in Equation 6). The decrease in prospective motor error is much slower than under LQR (Figure 4A, bottom), and is non-selective, with errors along potent and null directions being eliminated at the same rate (compare Figure 4F, top and bottom).

### Preparatory control in other M1 models

So far we have shown that feedback control is essential for rapid movement preparation in a specific model architecture. The inhibition-stabilized network (ISN) model we have used (Figure 2A and Figure 5A) has strong internal dynamics, whose “nonnormal” nature (Figure 5E; Trefethen and Embree, 2005) gives rise to pronounced transient amplification of a large subspace of initial conditions (Hennequin et al., 2014). This raises the concern that this ISN network might be unduly high-dimensional and difficult to control. After all, feedback control might not be essential in other models of movement generation for which the naive feedforward strategy might be good enough. To address this concern, we implemented optimal control in two other classes of network models which we trained to produce eight straight reaches in the same way as we trained the ISN, by optimizing the readout weights and the initial conditions (Figure 5B and C; Section ⋆2.1.4). Importantly, we also optimized aspects of the recurrent connectivity: either all recurrent connections (‘full’ networks, Figure 5B), or a low-rank parameterization thereof (‘low-rank’ networks, Figure 5C; Sussillo and Abbott, 2009; Mastrogiuseppe and Ostojic, 2018). We trained 10 instances of each network class, and also produced 10 new ISN networks for comparison. Empirically, we found that training was substantially impaired by the addition of the condition-independent movement-epoch input ***h***(*t*) in Equation 2 in the full and low-rank networks. We therefore dispensed with this input in the training of all these networks, as well as in the newly-trained ISNs for fair comparison (the results presented below also hold for ISNs trained with ***h***(*t*) ≠ 0).

**Figure 5:**
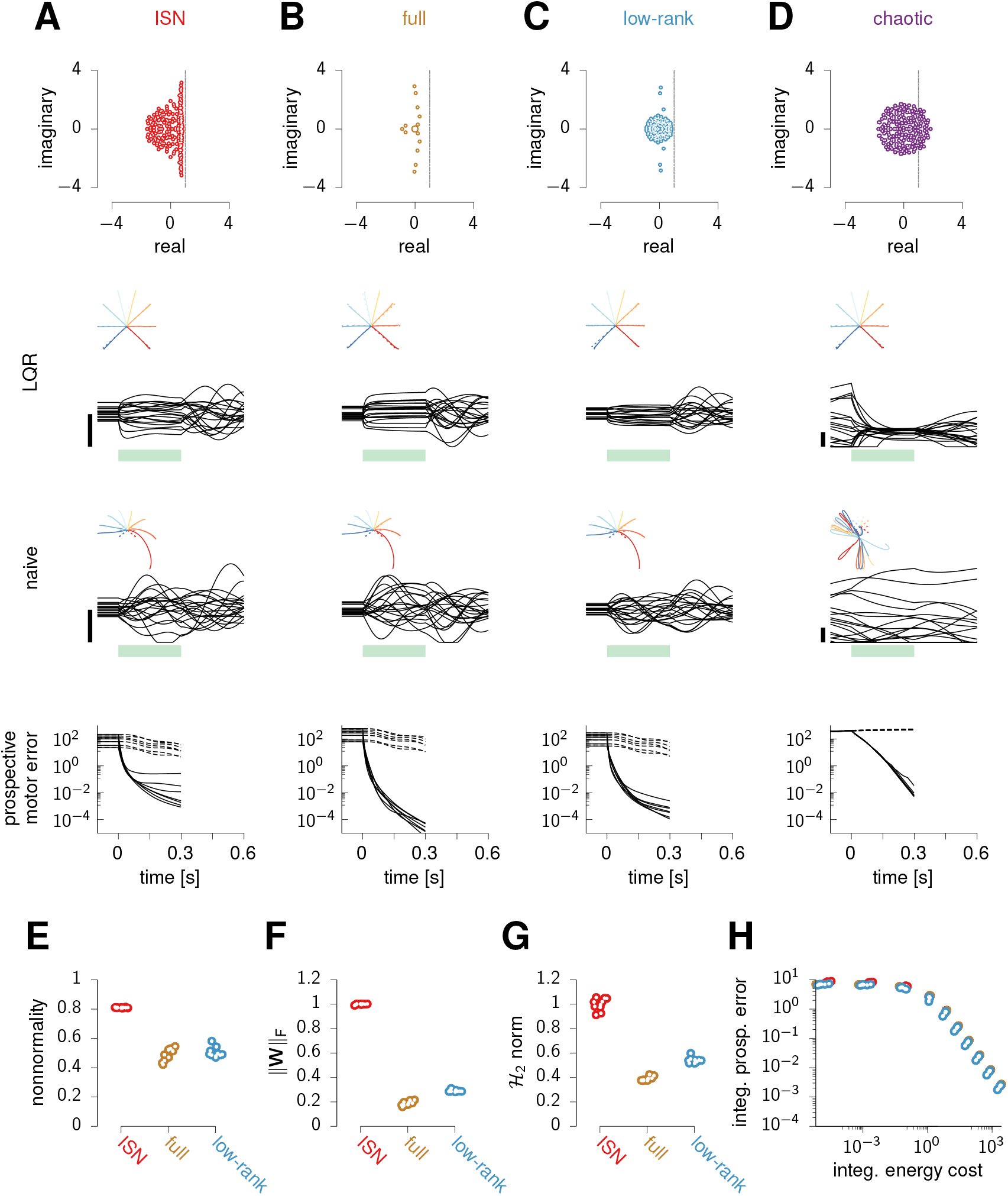
Optimal preparatory control benefits other models of movement generation. **(A)** ISN model. Top: eigenvalues of the connectivity matrix. Middle: activity of 20 example neurons during optimal (LQR) and naive feedforward preparatory control and subsequent execution of one movement, with hand trajectories shown as an inset for all movements. The green bar marks the preparatory period. Bottom: prospective motor error under LQR (solid) and naive feedforward (dashed) preparation, for each movement. **(B-D)** Same as (A), for a representative instance of each of the three other network classes (see text). **(E-G)** Index of nonnormality (E), Frobenius norm of the connectivity matrix ║***W***║_F_ (F), and 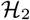 norm (G) for the 10 network instances of each class (excluding chaotic networks for which these quantities are either undefined or uninterpretable). Dots are randomly jittered horizontally for better visualization. Both ║***W***║_F_ and the 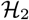 norm are normalized by the ISN average. **(H)** Quantification of controllability in the various networks. Optimal control cost (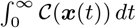 in Equation 6) against associated control energy cost 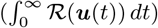, for different values of the energy penalty parameter λ, and for each network (same colors as E-G). Note that the naive feedforward strategy corresponds to the limit of zero energy cost (horizontal asymptote).

The full and low-rank networks successfully produce the correct hand trajectories after training (Figure 5B and C, middle), relying on dynamics with oscillatory components qualitatively similar to the ISN’s (eigenvalue spectra in Figure 5A-C, top). They capture essential aspects of movement-epoch population dynamics in monkey M1, to a similar degree as the ISN does (Figure S6). Nevertheless, the trained networks differ quantitatively from the ISN in ways that would seemingly make them easier to control. First, they are less non-normal (Figure 5E). Second, they have weaker internal dynamics than the ISN, as quantified by the average squared magnitude of their recurrent connections (Figure 5F). Third, these weaker dynamics translate into smaller impulse responses overall, as quantified by the 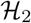 norm (Figure 5G; STAR Methods; Hennequin et al., 2014; Kao and Hennequin, 2019).

Although these quantitative differences suggest that optimal feedback control might be superfluous in the full and low-rank networks, we found that this is not the case. In order to achieve a set prospective control performance, all networks require the same total input energy (Figure 5H). In particular, the feedforward strategy (limit of zero input energy) performs equally badly in all networks: preparation is unrealistically slow, with 300 ms of preparation still resulting in large reach distortions (Figure 5A-C, bottom; recall also Figure 2B). In fact, we were able to show formally that the performance of feedforward control depends only on the target torques specified by the task, but not on the details of how these targets are achieved through specific initial conditions, recurrent connectivity ***W***, and readout matrix ***C*** (Supplementary Math Note 2). Stronger still, our derivations explain why the movement errors resulting from insuffiently long feedforward preparation are identical in every detail across all networks (Figure 5A-C, bottom).

Finally, we also considered networks in the chaotic regime with fixed and strong random connection weights and the same threshold-linear activation function (‘chaotic’ networks, Figure 5D; Kadmon and Sompolinsky, 2015; Mastrogiuseppe and Ostojic, 2017). In these networks, since static inputs are unable to quench chaos, the naive feedforward strategy cannot even establish a fixed point, let alone a correct one (Figure 5D, bottom). However, by adapting the optimal feedback control solution to the nonlinear case (STAR Methods), we found that it successfully quenches chaos during preparation and enables fast movement initiation (Figure 5D, middle).

In summary, the need for feedback control during preparation is not specific to our particular ISN model, but emerges broadly in networks of various types trained to produce reaches.

### Orthogonal preparatory and movement subspaces

Optimal anticipatory control also accounts for a prominent feature of monkey motor cortex responses during motor preparation and execution: across time and reach conditions, activity spans orthogonal subspaces during the two epochs. To show this, we followed Elsayed et al. and performed principal component analysis (PCA) on trial-averaged activity during the two epochs separately, in both the ISN model of Figure 4 and the two monkey datasets (Figure 6A). We then examined the fraction of variance explained by both sets of principal components (prep-PCs and move-PCs) during each epoch. Activity in the preparatory and movement epochs are, respectively, approximately 4- and 6-dimensional for the model, 5- and 7-dimensional for monkey J, and 8- and 7-dimensional for monkey N (assessed by the “participation ratio”). Moreover, consistent with the monkey data, prep-PCs in the model account for most of the activity variance during preparation (by construction; Figure 6B, left), but account for little variance during movement (Figure 6B, right). Similarly, move-PCs capture little of the preparatory-epoch activity variance.

**Figure 6:**
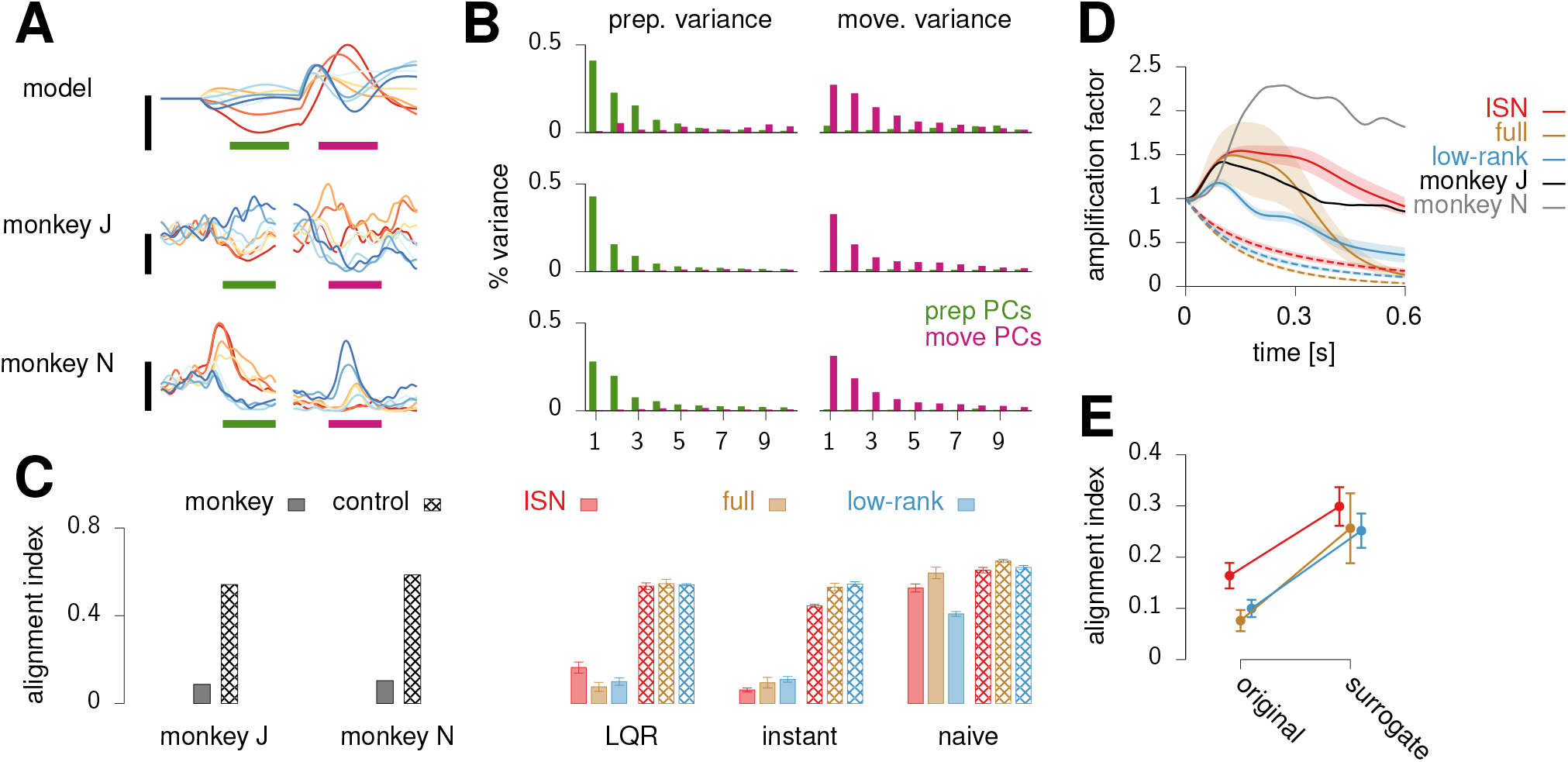
Reorganization between preparatory and movement activity in model and monkey. **(A)** Example single-neuron PSTHs in model (top) and monkey M1/PMd (middle and bottom), for each reach (c.f. Figure 2B). **(B)** Fraction of variance (time and conditions) explained during movement preparation (left) and execution (right) by principal components calculated from preparatory (green) and movement-related (magenta) trial-averaged activity. Only the first 10 components are shown for each epoch. Variance is across reach conditions and time in 300 ms prep. and move. windows indicated by green and magenta bars in (A). The three rows correspond to those of (A). **(C)** Alignment index calculated as in Elsayed et al. (2016), for the two monkeys (left) and the three classes of trained networks (colors as in Figure 5A-C) under three different preparation strategies (LQR, instant, naive; right). Here, preparation is long enough (500 ms) that even the naive feedforward strategy leads to the correct movements in all networks. Hashed bars show the average alignment index between random-but-constrained subspaces drawn as in Elsayed et al. (2016). **(D)** Amplification factor, quantifying the growth of the centered population activity vector ***x***(*t*) – ***x***_sp_ during the course of movement, relative to the pre-movement state (STAR Methods). It is shown here for the two monkeys (black and gray), as well as for the three classes of trained networks (solid) and their surrogate counterparts (dashed). Shaded region denotes ±1 s.t.d. around the mean across the 10 instances of each network class. To isolate the autonomous part of the movement-epoch dynamics, here we set ***h***(*t*) = 0 in Equation 2. **(E)** Alignment index under the ‘instant’ preparation strategy, for the original trained networks and their surrogates. Error bars denote ±1 s.t.d. across the 10 networks of each class.

To systematically quantify this (lack of) overlap for all the trained models of Figure 5A-C and compare with monkey data, we used Elsayed et al.’s “alignment index”. This is defined as the amount of preparatoryepoch activity variance captured by the top *K* move-PCs, further normalized by the maximum amount of variance that any *K*-dimensional subspace can capture. Here, *K* was chosen such that the top *K* prep-PCs capture 80% of activity variance during the preparatoryepoch. Both monkeys have a low alignment index (Figure 6C, left), much lower than a baseline expectation reflecting the neural activity covariance across both task epochs (“random” control in Elsayed et al., 2016). All trained models show the same effect under optimal preparatory control (Figure 6C, ‘LQR’). Importantly, alignment indices arising from the naive feedforward solution are much higher and close to the random control.

To understand the origin of such orthogonality, we first note that the preparatory end-states themselves tend to be orthogonal to movement-epoch activity in the models. Indeed, artificially clamping network activity to its end state during the whole preparatory epoch yields low alignment indices in all trained networks (Figure 6C, ‘instant’). We hypothesize that this effect arises from the fact that, as in monkeys, our trained networks all transiently amplify the initial condition at movement onset (Figure 6D). Since the autonomous (near-linear) dynamics that drive these transients are stable, the initial growth of activity at movement onset must be accompanied by a rotation away from the initial condition. Otherwise the growth would continue, contradicting stability. To substantiate this interpretation, for each network that we trained, we built a surrogate network that achieved the same task without resorting to transient growth (STAR Methods). We thus predicted higher alignment indices in these surrogate networks. To construct them, we noted that by applying a similarity transformation simultaneously to ***W, C*** and the initial conditions, one can arbitrarily suppress transient activity growth at movement onset yet maintain performance in the task in the linear regime (a similarity transformation does not change the overall transfer function from initial condition to output torques). In particular, the similarity transformation that diagonalizes ***W*** returns networks for which the magnitude of movement-epoch population activity can only decay during the course of movement, unlike that of the trained networks (Figure 6D, dashed). Consistent with our hypothesis, these surrogate networks have a higher ‘instant’ alignment index (Figure 6F).

So far, we have shown that orthogonality between *late*-preparatory and movement subspaces emerges generically in regularized trained networks. However, this does not fully explain why the alignment index is low under LQR in all models when early-/mid-preparatory activity is considered instead, as in Elsayed et al. (2016). In fact, under the naive feedforward strategy, early preparatory activity can be shown to be exactly the negative image of movement-epoch activity, up to a relatively small constant offset (Supplementary Math Note 3). Thus, the temporal variations of early preparatory- and movement-epoch activity occur in a shared subspace, generically yielding a high alignment index (Figure 6C, ‘naive’). In contrast, optimal control rapidly eliminates preparatory errors along potent directions, which contribute significantly to movement-epoch activity: if ***x***(*t*) during movement had no potent component, by definition the movement would stop immediately. Thus, one expects optimal control to remove a substantial fraction of overlap between preparatory- and movement-epoch activity. What is more, by quenching temporal fluctuations along potent directions early during preparation, optimal feedback also suppresses the subsequent transient growth of activity that these potent fluctuations would have normally produced under the naive strategy (and which are indeed produced during movement). The combination of these primary and secondary suppressive effects results in the lower alignment index that we observe (Figure 6C, ‘LQR’).

In summary, orthogonality between preparatory and movement activity in monkey M1 is consistent with optimal feedback control theory. In the ISN, as well as in other architectures trained on the same task, orthogonality arises robustly from optimal preparatory control, but not from feedforward control.

### Circuit model for preparatory control: a gated thalamocortical loop

So far we have not discussed the source of optimal preparatory inputs ***u***(*t*), other than saying that they close a feedback loop from the cortex onto itself (Equation 7). Such a loop could in principle be absorbed in local modifications of recurrent cortical connectivity (Sussillo and Abbott, 2009). However, we are not aware of any mechanism that could implement nearinstant on/off switching of a select subset of recurrent synapses, as required by the model at onset of preparation (ON) and movement (OFF). If, instead, the preparatory loop were to pass through another brain area, fast modulation of excitability in that relay area would provide a rapid and flexible switch (Ferguson and Cardin, 2020). We therefore propose the circuit model shown in Figure 7A, where the motor thalamus acts as a relay station for cortical feedback (Guo et al., 2017; Nakajima and Halassa, 2017). The loop is gated on/off at preparation onset/offset by the (dis)-inhibitory action of basal ganglia outputs (Jin and Costa, 2010; Cui et al., 2013; Dudman and Krakauer, 2016; Halassa and Acsaády, 2016; Logiaco et al., 2019). Specifically, cortical excitatory neurons project to 160 thalamic neurons, which make excitatory backprojections to a pool of 100 excitatory (E) and 100 inhibitory (I) neurons in cortex layer 4. In turn, these layer 4 neurons provide both excitation and inhibition to the main cortical network, thereby closing the control loop. Here, inhibition is necessary to capture the negative nature of optimal feedback. In addition to thalamic feedback, the cortical network also receives a movement-specific constant feedforward drive during preparation (analogous to ***u***⋆ in Equation 7 for the standard LQR algorithm; this could also come from the thalamus).

**Figure 7:**
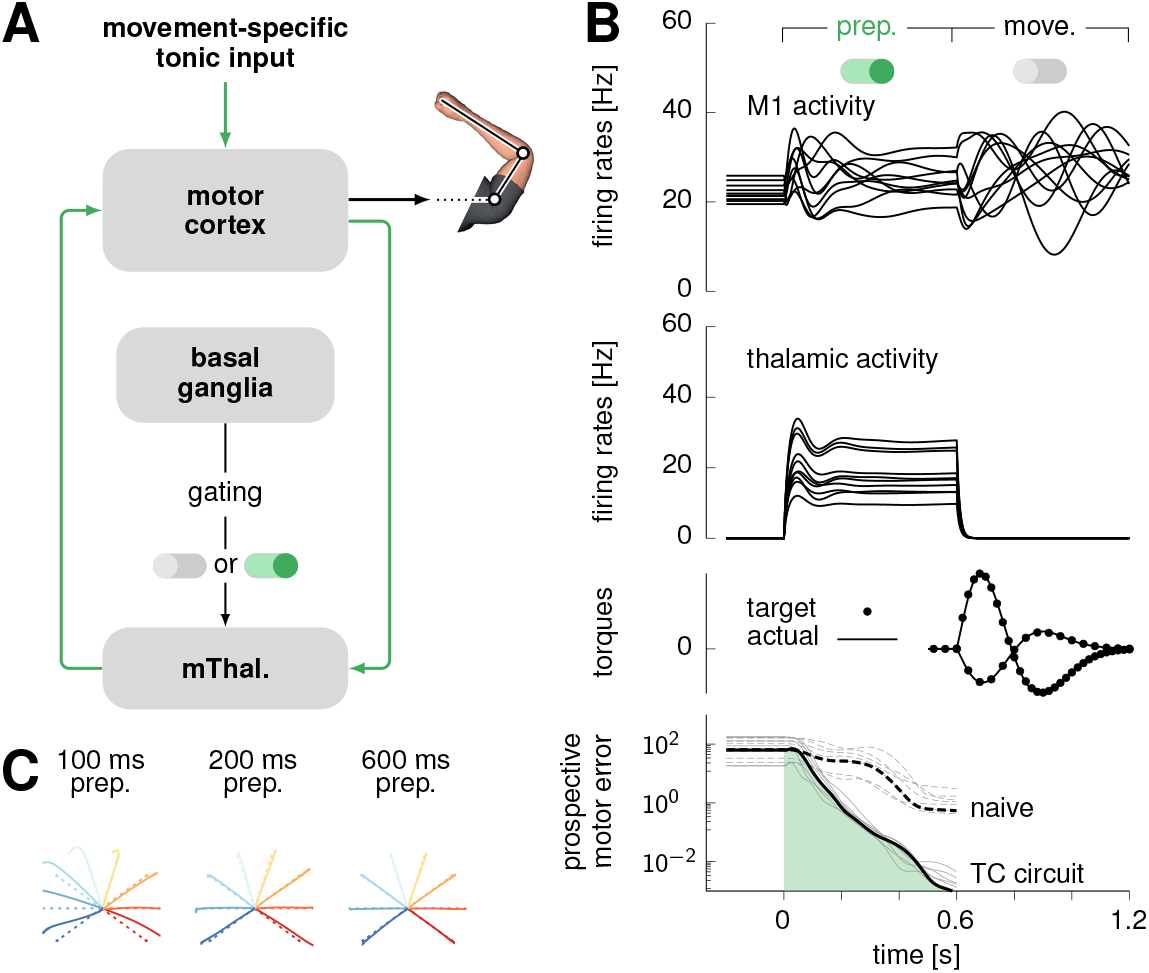
Optimal movement preparation via a gated thalamo-cortical loop. **(A)** Proposed circuit architecture for the optimal movement preparation (cf. text). **(B)** Cortical (top) and thalamic (upper middle) activity (10 example neurons), generated torques (lower middle), and prospective motor error (bottom) during the course of movement preparation and execution in the circuit architecture shown in (A). The prospective motor error for the naive strategy is shown as a dotted line as in Figure 4A. All black curves correspond to the same example movement (324-degree reach), and gray curves show the prospective motor error for the other 7 reaches. **(C)** Hand trajectories (solid) compared to target trajectories (dashed) for the eight reaches, triggered after 100 ms (left), 200 ms (middle) and 600 ms (right) of motor preparation.

The detailed patterns of synaptic efficacies in the thalamo-cortical loop are obtained by solving the same control problem as above, based on the minimization of the cost functional in Equation 6 (STAR Methods). Importantly, the solution must now take into account some key biological constraints: (i) feedback must be based on the activity of the cortical E neurons only, (ii) thalamic and layer-4 neurons have intrinsic dynamics that introduce lag, and (iii) the sign of each connection is constrained by the nature of the presynaptic neuron (E or I).

The circuit model we have obtained meets the above constraints and enables flexible, near-optimal anticipatory control of the reaching movements (Figure 7B). Before movement preparation, thalamic neurons are silenced due to strong inhibition from basal ganglia outputs (not explicitly modelled), keeping the thalamocortical loop open (inactive) by default. At the onset of movement preparation, rapid and sustained disinhibition of thalamic neurons restores responsiveness to cortical inputs, thereby closing the control loop (on/off switch in Figure 7B, top). This loop drives the cortical network into the appropriate preparatory subspace, rapidly reducing prospective motor errors as under optimal LQR feedback (Figure 7B, bottom). To trigger movement, the movement-specific tonic input to cortex is shut off, and the basal ganglia resume sustained inhibition of the thalamus. Thus, the loop re-opens, which sets off the uncontrolled dynamics of the cortical network from the right initial condition to produce the desired movement (Figure 7C).

The neural constraints placed on the feedback loop are a source of suboptimality with respect to the unconstrained LQR solution of Figure 4. Nevertheless, movement preparation by this thalamocortical circuit remains fast, on par with the shortest preparation times of primates in a quasi-automatic movement context (Lara et al., 2018) and much faster than the naive feedforward strategy (Figure 7B, bottom).

### Model prediction: selective elimination of preparatory errors following optogenetic perturbations

An essential property of optimal preparatory control is the selective elimination of preparatory errors along prospectively potent directions. As a direct corollary, the model predicts selective recovery of activity along those same directions following preparatory perturbations of the dynamics, consistent with results by Li et al. (2016) in the context of a delayed directional licking task in mice. Here, we spell out this prediction in the context of reaching movements in primates by simulating an experimental perturbation protocol similar to that of Li et al. (2016), and by applying their analysis of population responses. These concrete predictions should soon become testable with the advent of optogenetics techniques in primates (O’Shea et al., 2018).

As our E/I cortical circuit model operates in the inhibition-stabilized regime (Hennequin et al., 2014; Tsodyks et al., 1997; Ozeki et al., 2009; Sanzeni et al., 2019), we were able to use the same photoinhibition strategy as in Li et al. (2016) to silence the cortical network (Figure 8A) in the thalamo-cortical circuit of Figure 7. We provided strong excitatory input to a random subset (60%) of inhibitory neurons, for a duration of 400 ms starting 400 ms after preparation onset. We found that “photoinhibition” has mixed effects on the targeted neurons: some are caused to fire at higher rates, but many are paradoxically suppressed (Figure 8B, top). For E cells and untargeted I cells, though, the effect is uniformly suppressive, as shown in Figure 8B (middle and bottom).

**Figure 8:**
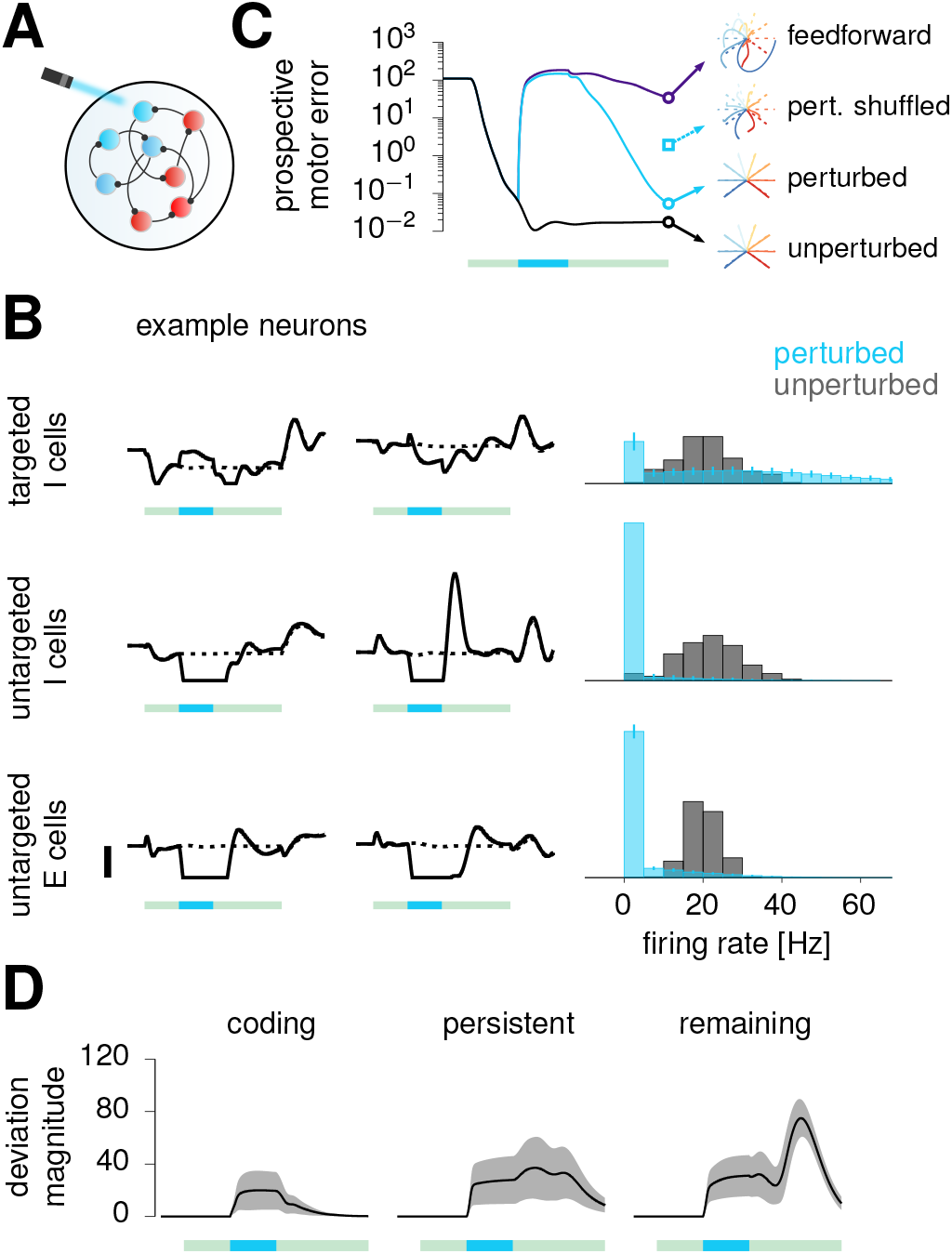
Testable prediction: selective recovery from preparatory perturbations. **(A)** Illustration of perturbation via “photoinhibition”: a subset (60%) of I neurons in the model are driven by strong positive input. **(B)** Left: firing rates (solid: perturbed; dashed: unperturbed) for a pair of targeted I cells (top), untargeted I cells (middle) and E cells (bottom). Green bars (1.6 s) mark the movement preparation epoch, and embedded turquoise bars (400 ms) denote the perturbation period. Right: histogram of firing rates observed at the end of the perturbation (turquoise), and at the same time in unperturbed trials (gray). Error bars show one standard deviation across 300 experiments, each with a different random set of targeted I cells. **(C)** Prospective motor error, averaged across movements and perturbation experiments, in perturbed (turquoise) vs. unperturbed (black) conditions. Subsequent hand trajectories are shown for one experiment of each condition (middle and bottom insets; dashed lines show target trajectories). These are compared with the reaches obtained by randomly shuffling final preparatory errors *δ**x*** across neurons, and re-simulating the cortical dynamics thereafter (turquoise square mark). The purple line shows the performance of an optimal *feedforward* strategy, which pre-computes the inputs that would be provided to the cortex under optimal feedback control, and subsequently replays those inputs at preparation onset without taking the state of the cortex into account anymore. **(D)** Magnitude of the deviation caused by the perturbation in the activity of the network projected into the coding subspace (left), the persistent subspace (center) and the remaining subspace (right). Lines denote the mean across perturbation experiments, and shadings indicate ±1 s.t.d. Green and turquoise bars as in (B).

The perturbation transiently resets the prospective motor error to pre-preparation level, thus nullifying the benefits of the first 400 ms of preparation (Figure 8C). Following the end of the perturbation, the prospective motor error decreases again, recovering to unperturbed levels (Figure 8C, solid vs. dashed) and thus enabling accurate movement production (compare middle and bottom hand trajectories). This is due to the selective elimination of preparatory errors discussed earlier (Figure 4F): indeed, shuffling *δ**x*** across neurons immediately prior to movement, thus uniformizing the distribution of errors in different state space directions, leads to impaired hand trajectories (Figure 8C, top right).

We next performed an analysis qualitatively similar to Li et al.‘s (STAR Methods). We identified a “coding subspace” (CS) that accounts for most of the across-condition variance in firing rates towards the end of movement preparation in unperturbed trials. Similarly, we identified a “persistent” subspace (PS) that captures most of the activity difference between perturbed and unperturbed trials towards the end of preparation, regardless of the reach direction. Finally, we also contructed a third subspace, constrained to be orthogonal to the PS and the CS, and capturing most of the remaining variance across the different reaches and perturbation conditions (“remaining subspace”, RS).

We found the CS and the PS to be nearly orthogonal (minimum subspace angle of 89 degrees) even though they were not constrained to be so. Moreover, the perturbation causes cortical activity to transiently deviate from unperturbed trajectories nearly equally in each of the three subspaces (Figure 8D). Remarkably, however, activity recovers promptly along the CS, but not along the other two modes. In fact, the perturbation even grows transiently along the PS during early recovery. Such selective recovery can be understood from optimal preparation eliminating errors along directions with motor consequences, but (owing to energy constraints) not in other inconsequential modes. Indeed, the CS is by definition a prospectively potent subspace: its contribution to the preparatory state is what determines the direction of the upcoming reach. In contrast, the PS and the RS are approximately prospectively null (respectively 729 times and 8 times less potent than the CS, by our measure 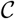 of motor potency in Figure 3D). This not only explains why the CS and PS are found to be near-orthogonal, but also why the thalamocortical dynamics implementing optimal preparatory control only quench perturbation-induced deviations in the CS.

In summary, optogenetic perturbations of M1 could be used to test a core prediction of optimal preparatory control, namely the selective recovery of activity in subspaces that carry movement information, but not in others.

Finally, perturbation experiments can also help distinguish between optimal feedback control and feedforward control strategies. The naive feedforward strategy discussed previously can already be ruled out for being unrealistically slow. However, it is easy to conceive of other feedforward control mechanisms that would be more difficult to distinguish from feedback control. In particular, consider “optimal” feedforward control, which pre-computes the optimal feedback inputs once and for all, and provides them each time the same movement is to be prepared. By definition, this feedforward strategy is indistiguishable from optimal feedback control in the absence of noise or perturbation during preparation (compare black and purple lines in Figure 8C before the onset of perturbation). However, unlike optimal feedback, the optimal feedforward strategy cannot react to state perturbations during preparation as its inputs are pre-computed and thus unable to adapt to unexpected changes in the cortical state (compare purple and turquoise lines in Figure 8C).

## Discussion

Neural population activity in cortex can be accurately described as arising from low-dimensional dynamics (Churchland et al., 2012; Mante et al., 2013; Carnevale et al., 2015; Seely et al., 2016; Barak, 2017; Cunningham and Byron, 2014; Michaels et al., 2016). These dynamics unfold from a specific initial condition on each trial, and indeed these “preparatory states” predict the subsequent evolution of both neural activity and behaviour in single trials of the task (Churchland et al., 2010; Pandarinath et al., 2018; Remington et al., 2018; Sohn et al., 2019). In addition, motor learning may rely on these preparatory states partitioning the space of subsequent movements (Sheahan et al., 2016).

How are appropriate initial conditions reached in the first place? Here, we have formalized movement preparation as an optimal control problem, showing how to translate anticipated motor costs phrased in terms of muscle kinematics into costs on neural activity in M1. Optimal preparation minimizes these costs, and the solution is feedback control: the cortical network must provide corrective feedback to itself, based on prospective motor errors associated with its current state. In other words, optimal preparation may rely on an implicit forward model predicting the future motor consequences of *preparatory activity*—not motor commands, as in classical theories (Wolpert et al., 1995; Desmurget and Grafton, 2000; Scott, 2012)—and feeding back these predictions for online correction of the cortical trajectory. Thus, where previous work has considered the motor cortex as a controller of the musculature (Todorov, 2000; Lillicrap and Scott, 2013; Sussillo et al., 2015), our work considers M1 and the body jointly as a compound system under preparatory control by other brain areas.

One of the key assumptions of this work is that the cortical dynamics responsible for reaching are at least *partially* determined by the inital state (we do not assume that they are autonomous). If, on the contrary, M1 activity were purely input-driven, there would be no need for preparatory control, let alone an optimal one. On the one hand, perturbation experiments in mice suggest thalamic inputs are necessary for continued movement generation and account for a sizeable fraction (though not all) of cortical activity during movement production (Sauerbrei et al., 2020). On the other hand, experiments in non-human primates provide ample evidence that preparation does occur, irrespective of context (Lara et al., 2018), and that initial pre-movement states influence subsequent behaviour (Churchland et al., 2010; Afshar et al., 2011; Shenoy et al., 2013). Thus, preparatory control is likely essential for accurate movement production.

### Sloppy preparation for accurate movements

A core insight of our analysis is that preparatory activity must be constrained in a potent subspace impacting future motor output, but is otherwise free to fluctuate in a nullspace. This readily explains two distinctive features of preparatory activity in reaching monkeys: (i) that pre-movement activity on zero-delay trials needs not reach the state achieved for long movement delays (Ames et al., 2014), and (ii) that nevertheless movement is systematically preceded by activity in the same preparatory subspace irrespective of whether the reach is self-initiated, artificially delayed, or reactive and fast (Lara et al., 2018). In our model, preparatory activity converges rapidly in the subspace that matters, such that irrespective of the delay (above 50 ms), preparatory activity is always found to have some component in this common subspace as in Lara et al. (2018). Moreover, exactly which of the many acceptable initial conditions is reached by the end of the delay depends on the delay duration. Thus, our model predicts that different preparatory end-states will be achieved for short and long delays, consistent with the results of Ames et al. (2014). Finally, these end-states also depends on the activity prior to preparation onset. This would explain why Ames et al. (2014) observed different pre-movement activity states when preparation started from scratch and when it was initiated by a change in target that interrupted a previous preparatory process. In the same vein, Sauerbrei et al. (2020) silenced thalamic input to mouse M1 early during movement, and observed that M1 activity did not recover to the pre-movement activity seen in unperturbed trials, even when the movement was successfully performed following the perturbation. This result is expected in our model if the perturbation is followed by a new preparatory phase in which the effect of the perturbation rapidly vanishes along prospectively potent dimensions, but subsists in the nullspace.

### Preparing without moving

How can preparatory M1 activity not cause premature movement, if M1 directly drives muscles? Kaufman et al. (2014) argued that cortical activity may evolve in a coordinated way at the population level so as to remain in the nullspace of the muscle readout. Here, we have not explicitly penalized premature movements as part of our control objective (Equation 6). As a result, while preparatory activity enters the nullspace eventually (because we constrained the movement-seeding initial conditions to belong to it), optimal preparatory control yields small activity excursions outside the nullspace early during preparation (Figure S7), causing the hand to drift absent any further gating mechanism. Nevertheless, one can readily penalize such premature movements in our framework and obtain network models that prepare rapidly “in the nullspace” via closed-loop feedback (Figure S7; STAR Methods). Importantly, a naive feedforward control strategy is unable to contain the growth of movement-inducing activity during preparation.

### Thalamic control of cortical dynamics

The mathematical structure of the optimal control solution suggested a circuit model based on cortico-cortical feedback. We have proposed that optimal feedback can be implemented as a cortico-thalamo-cortical loop, switched ON during movement preparation and OFF again at movement onset. The ON-switch occurs through fast disinhibition of those thalamic neurons that are part of the loop. Our model thus predicts a large degree of specificity in the synaptic interactions between cortex and thalamus (Halassa and Sherman, 2019; Huo et al., 2020), as well as a causal involvement of the thalamus in movement preparation (Guo et al., 2017; Sauerbrei et al., 2020). Furthermore, the dynamical entrainment of thalamus with cortex predicts tuning of thalamic neurons to task variables, consistent with a growing body of work showing specificity in thalamic responses (Nakajima and Halassa, 2017; Guo et al., 2017; Rikhye et al., 2018). For example, we predict that neurons in the motor thalamus should be tuned to movement properties, for much the same reasons that cortical neurons are (Todorov, 2000; Lillicrap and Scott, 2013; Omrani et al., 2017). Finally, we speculate that an on/off switch on the thalamocortical loop is provided by one of the output nuclei of the basal ganglia (SNR or GPi), or a subset of neurons therein. Thus, exciting these midbrain neurons during preparation should prevent the thalamus from providing the necessary feedback to the motor cortex, thereby slowing down the preparatory process and presumably increasing reaction times. In contrast, silencing these neurons during movement should close the preparatory thalamocortical loop at a time when thalamus would normally disengage. This should modify effective connectivity in the movement-generating cortical network and impair the ongoing movement. These are predictions that could be tested in future experiments.

Thalamic control of cortical dynamics offers an attractive way of performing nonlinear computations (Sussillo and Abbott, 2009; Logiaco et al., 2019). Although both preparatory and movement-related dynamics are approximately linear in our model, the transition from one to the other (orchestrated by the basal ganglia) is highly nonlinear. Indeed, our model can be thought of as a switching linear dynamical system (Linderman et al., 2017). Moreover, gated thalamocortical loops are a special example of achieving nonlinear effects through gain modulation. Here, it is the thalamic population only that is subjected to abrupt and binary gain modulation, but changes in gain could also affect cortical neurons. This was proposed recently as a way of expanding the dynamical repertoire of a cortical network (Stroud et al., 2018).

Switch-like nonlinearities may have relevance beyond movement preparation, e.g. for movement execution. In our model, different movement patterns are produced by different initial conditions seeding the same generator dynamics. However, we could equally well have generated each reach using a different movement-epoch thalamocortical loop. Logiaco et al. have recently explored this possibility, showing that gated thalamocortical loops provide an ideal substrate for flexible sequencing of multiple movements. In their model, each movement is achieved by its own loop (involving a shared cortical network), and the basal ganglia orchestrate a chain of thalamic disinhibitory events, each spatially targeted to activate those neurons that are responsible for the next loop in the sequence (Logiaco et al., 2019). Interestingly, their cortical network must still be properly initialized prior to each movement chunk, as it must in our model. For this, they proposed a generic preparatory loop similar to the one we have studied here. However, theirs does not take into account the degeneracies in preparatory states induced by prospective motor costs, which ours exploits. In sum, our model and theirs address complementary facets of motor control (preparation and sequencing), and could be combined into a single model.

To conclude, we have proposed a new theory of movement preparation and studied its implications for various models of movement-generating dynamics in M1. While these models capture several salient features of movement-epoch activity, they could be replaced by more accurate, data-driven models (Pandarinath et al., 2018) in future work. This would enable our theory to make detailed quantitative predictions of preparatory activity, which could be tested further in combination with targeted perturbation experiments. Our control-theoretic framework could help elucidate the role of the numerous brain areas that collectively control movement (Svoboda and Li, 2018), and make sense of their hierarchical organization in nested loops.

## Acknowledgments

This work was supported by a Seed Award from the Wellcome Trust (202111/Z/16/Z), a Trinity-Henry Barlow scholarship (T-C.K.), and a scholarship from the Ministry of Education, ROC Taiwan (T-C.K.). We are grateful to Mark Churchland, Matthew Kaufman and Krishna Shenoy for sharing their data and for discussions, to Karel Svoboda and his lab for hospitality and discussions, to James Heald for sharing the 3D character drawn in Figure 1A, and to Marine Schimel and Kristopher Jensen for feedback on the manuscript.

## STAR⋆METHODS

### ⋆1 Resource availability

#### ⋆1.1 Lead contact

Further information and requests for resources and reagents should be directed to and will be fulfilled by the Lead Contact, Guillaume Hennequin.

#### ⋆1.2 Materials availability

This study did not generate new unique reagents.

#### ⋆1.3 Data and code availability

The full datasets have not been deposited in a public repository because they are made available to us by Krishna Shenoy, Mark Churchland, and Matt Kaufman. Requests for these recordings should be directed to them. The code generated during this study is available at https://github.com/hennequin-lab/optimal-preparation.

### ⋆2 Method details

#### ⋆2.1 A model for movement generation by cortical dynamics

##### ⋆2.1.1 Network dynamics

We model M1 as a network with two separate populations of *N*_E_ = 160 excitatory (E) neurons and *N*_I_ = 40 inhibitory (I) neurons, operating in the inhibition-stabilized regime (Tsodyks et al., 1997; Ozeki et al., 2009; Hennequin et al., 2014). We constructed its synaptic architecture using the algorithm we have previously described in Hennequin et al. (2014). Briefly, we iteratively updated the inhibitory synapses of a random network, with an initial spectral abscissa of 1.2, to minimize a measure of robust network stability. We implemented early stopping, terminating the stabilization procedure as soon as the spectral abscissa of the connectivity matrix dropped below −0.8.

We describe the dynamics of these *N* = *N*_E_ + *N*_I_ neurons by a standard nonlinear rate equation. Specifically, the vector 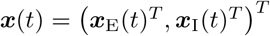 of internal neuronal “activations” obeys:

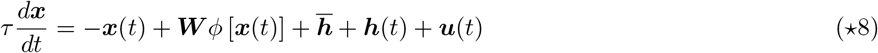

where *τ* is the single-neuron time constant, ***W*** is the synaptic connectivity matrix, and *ϕ*(*x*) = max(*x*, 0) is a static, rectified-linear nonlinearity – applied elementwise to ***x*** – that converts internal activation into momentary firing rates. The input consists of three terms: an input 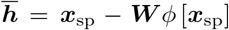 held constant throughout all phases of the task to instate a heterogeneous set of spontaneous firing rates ***x***_sp_ (elements drawn i.i.d. from 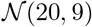); a transient, movement-condition-independent and spatially uniform α-shaped input bump

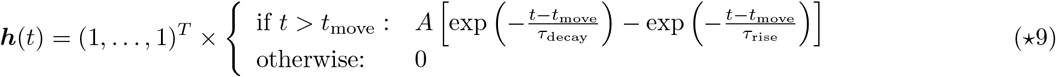

kicking in at movement onset (Kaufman et al., 2016); and a preparatory control input ***u***(*t*) (further specified below) whose role is to drive the circuit into a preparatory state appropriate for each movement.

We assume that the uncontrolled dynamics (***u*** = **0**) of this network directly drives movement. To actuate the two-link arm model described in the next section, the activity of the network is read out into two joint torques:

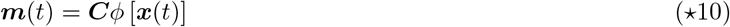

where 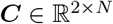 is such that its last *N*_I_ columns are zero, i.e. only the excitatory neurons contribute directly to the motor output. Although our simulations show that the muscle readouts ***m***(*t*) are very small during preparation, they do cause drift in the hand prior to movement onset (and therefore wrong movements afterwards) as they are effectively integrated twice by the dynamics of the arm (see below and Figure S7). For this reason, we artificially set ***m*** to zero during movement preparation.

##### ⋆2.1.2 Arm model

To simulate reaching movements, we used the planar two-link arm model previously described in Li and Todorov (2004), with parameters listed in Table ⋆1. The upper arm and the lower arm are connected at the elbow (Figure S1). The two links have lengths *L*_1_ and *L*_2_, masses *M*_1_ and *M*_2_, and moments of inertia *I*_1_ and *I*_2_ respectively. The lower arm’s center of mass is located at a distance *D*_2_ from the elbow. By considering the geometry of the upper and lower limb, we can write down the position of the hand as a vector **y**(*t*) given by

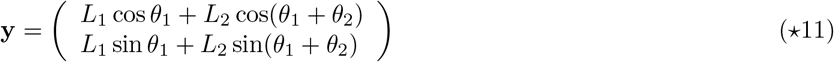

where the angles *θ*_1_ and *θ*_2_ are defined in Figure S1A. The joint angles ***θ*** = (*θ*_1_; *θ*_2_)^*T*^ evolve dynamically according to the differential equation

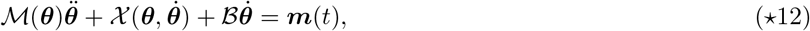

where ***m***(*t*) is the momentary torque vector (the output of the neural network, c.f. Equation ⋆10), 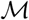 is the matrix of inertia, 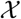 accounts for the centripetal and Coriolis forces, and 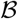 is a damping matrix representing joint friction.

These parameters are given by

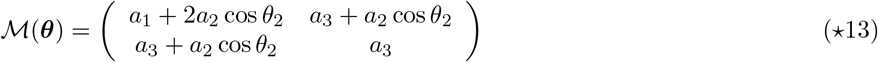

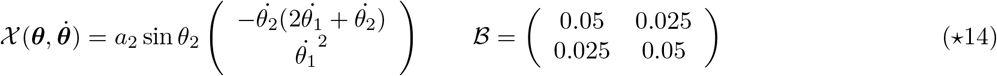

with 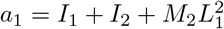, *a*_2_ = *M*_2_*L*_1_ *D*_2_, and *a*_3_ = *I*_2_.

**Table ⋆1:**
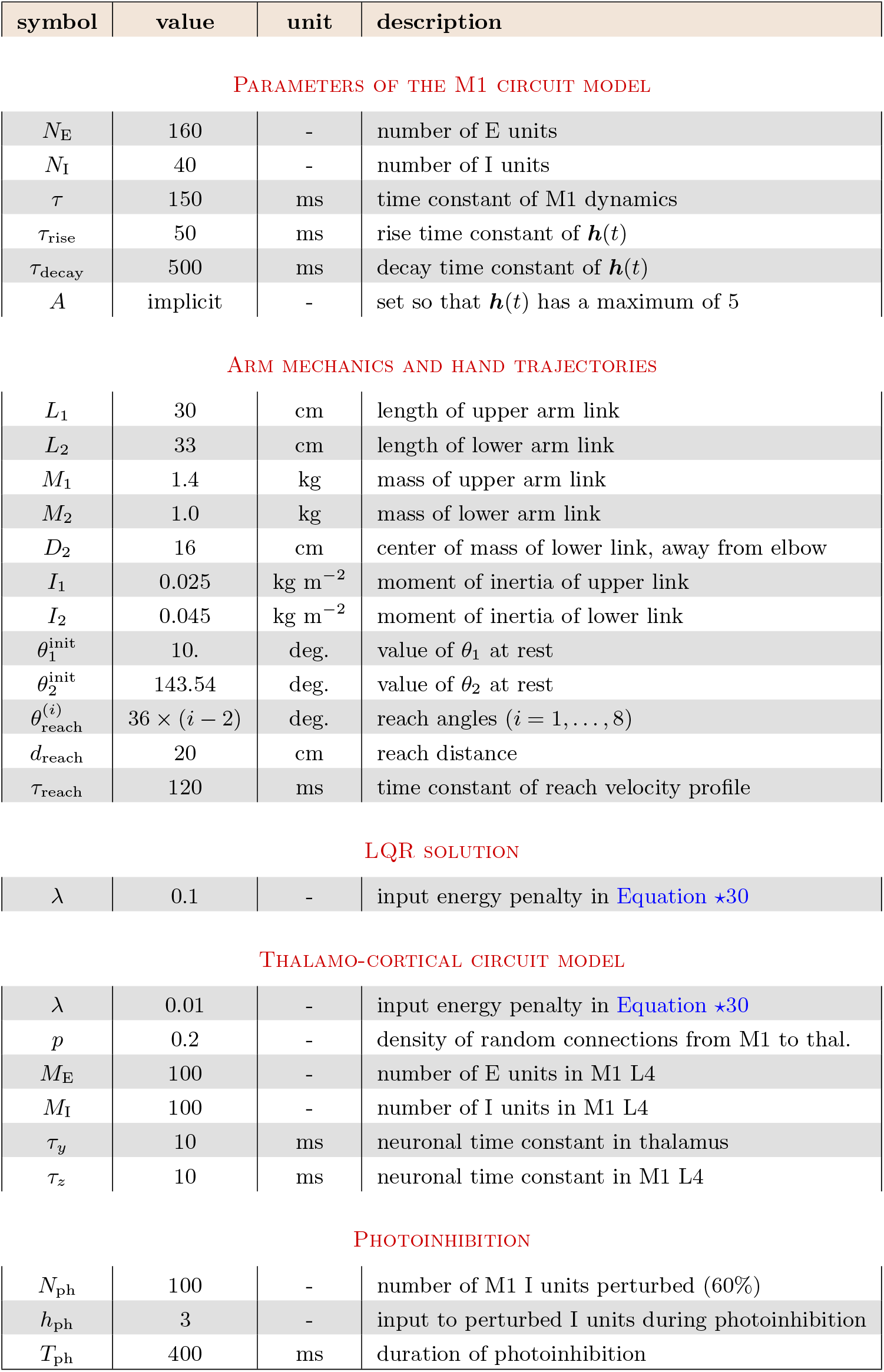
Generic parameters used throughout all simulations

The values of all the parameters used in this study are listed in the table below.
| symbol | value | unit | description |
| --- | --- | --- | --- |
| $N_E$ | 160 | - | number of E units |
| $N_I$ | 40 | - | number of I units |
| $\tau$ | 150 | ms | time constant of M1 dynamics |
| $\tau_{\text{rise}}$ | 50 | ms | rise time constant of $\mathbf{h}(t)$ |
| $\tau_{\text{decay}}$ | 500 | ms | decay time constant of $\mathbf{h}(t)$ |
| $A$ | implicit | - | set so that $\mathbf{h}(t)$ has a maximum of 5 |
| ARM MECHANICS AND HAND TRAJECTORIES |  |  |  |
| $L_1$ | 30 | cm | length of upper arm link |
| $L_2$ | 33 | cm | length of lower arm link |
| $M_1$ | 1.4 | kg | mass of upper arm link |
| $M_2$ | 1.0 | kg | mass of lower arm link |
| $D_2$ | 16 | cm | center of mass of lower link, away from elbow |
| $I_1$ | 0.025 | kg m<sup>-2</sup> | moment of inertia of upper link |
| $I_2$ | 0.045 | kg m<sup>-2</sup> | moment of inertia of lower link |
| $\theta_1^{\text{init}}$ | 10. | deg. | value of $\theta_1$ at rest |
| $\theta_2^{\text{init}}$ | 143.54 | deg. | value of $\theta_2$ at rest |
| $\theta_{\text{reach}}^{(i)}$ | $36 \times (i - 2)$ | deg. | reach angles ($i = 1, \dots, 8$) |
| $d_{\text{reach}}$ | 20 | cm | reach distance |
| $\tau_{\text{reach}}$ | 120 | ms | time constant of reach velocity profile |
| LQR SOLUTION |  |  |  |
| $\lambda$ | 0.1 | - | input energy penalty in Equation *30 |
| $\lambda$ | 0.01 | - | input energy penalty in Equation *30 |
| $p$ | 0.2 | - | density of random connections from M1 to thal. |
| $M_E$ | 100 | - | number of E units in M1 L4 |
| $M_I$ | 100 | - | number of I units in M1 L4 |
| $\tau_y$ | 10 | ms | neuronal time constant in thalamus |
| $\tau_z$ | 10 | ms | neuronal time constant in M1 L4 |
| PHOTOINHIBITION |  |  |  |
| $N_{\text{ph}}$ | 100 | - | number of M1 I units perturbed (60%) |
| $h_{\text{ph}}$ | 3 | - | input to perturbed I units during photoinhibition |
| $T_{\text{ph}}$ | 400 | ms | duration of photoinhibition |

##### ⋆2.1.3 Target hand trajectories and initial setup

We generated a set of eight target hand trajectories, namely straight reaches of length *d* = 20 cm going from the origin ((0, *d*) from the shoulder) into eight different directions, with a common bell-shaped scalar speed profile

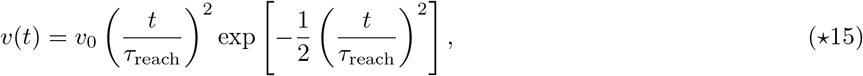

where *υ*_0_ is adjusted such that the hand reaches the target. Given these target hand trajectories, we solved for the required timecourse of the torque vector ***m***(*t*) through optimization, by backpropagating through the equations of motion of the arm (discretized using Euler’s method) to minimize the squared difference between actual and desired hand trajectories. We forced the initial torques at *t* = 0 to be zero, and also included a roughness penalty in the form of average squared torque gradient.

To calibrate the network for the production of the desired movement-specific torques 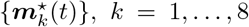, we optimized a set of eight initial conditions 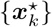, as well as the readout matrix ***C***, by minimizing the loss function:

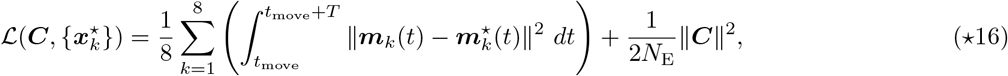

where *T* =1 second and ***m***_*k*_(*t*) depends on ***C*** and 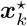 implicitly through the dynamics of the network. The first term of 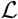 minimizes the squared difference between the actual and desired torque trajectories, while the second term penalizes ***C***’s squared Frobenius norm. We performed the optimization using the L-BFGS algorithm (Liu and Nocedal, 1989) and backpropagating through Equations ⋆8 and ⋆10. In addition, we parameterized the readout matrix ***C*** in such a way that its nullspace automatically contains both the spontaneous activity vector ***x***_sp_ and the movement-specific initial conditions 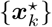, *k* = 1,…, 8. This is to ensure that (i) there is no muscle output during spontaneous activity and (ii) the network does not unduly generate muscle output at the end of preparation, before movement. More specifically, prior to movement preparation and long enough after movement execution, the cortical state is in spontaneous activity ***x***_sp_. By ensuring that ***C**ϕ*(***x***_sp_) = ***Cx***_sp_ = **0**, we ensure that our model network does not elicit movement “spontaneously”. Similarly, control inputs drive the cortical state ***x*** towards 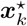, which it will eventually reach late in the preparation epoch; therefore, if we did not constrain 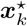 to be in ***C***’s null-space, premature movements would be elicited towards the end of preparation.

##### ⋆2.1.4 Training of other network classes

In Figures 5 and 6, we considered three other network types: ‘full’, ‘low-rank’, and ‘chaotic’ networks, which differed from the ISN in the way we constructed their synaptic connectivity matrices. We implemented 10 independent instances of each class.

For the chaotic networks, synaptic weights were drawn from a normal distribution with mean −25/*N* and variance 1.8^2^/*N*; this places the networks in the chaotic regime with a threshold-linear nonlinearity (Kadmon and Sompolinsky, 2015). We calibrated the chaotic networks for generating the desired hand trajectories by optimizing ***C*** and 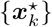 as described above.

Unlike the ISNs and the chaotic networks, we optimized the connectivity matrix ***W*** of the full and low-rank networks in addition to 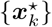 and ***C***, when we calibrated these networks for movement production. While we optimized every element of ***W*** for the full networks, we parameterized the low-rank networks as

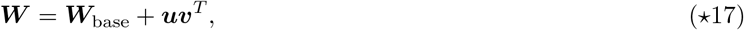

where ***uv***^*T*^ is a rank-5 perturbation to a random connectivity matrix ***W***_base_. Here, both ***u*** and 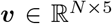 are free parameters, whereas ***W***_base_ is a fixed matrix with elements drawn anew for each network instance from 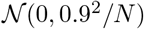. Following Sussillo et al. (2015), we regularized the ‘full’ and ‘low-rank’ networks by augmenting the loss function (Equation ⋆16) with an additional term: ║***W***║^2^/*N*^2^ for the ‘full’ networks and (║***u***║^2^ + ║***υ***║^2^)/5*N* for the ‘low-rank’ networks.

We found that the addition of the reach-independent input ***h***(*t*) during the movement epoch (Equation 2) made the ‘full’, ‘chaotic’, and ‘low-rank’ networks difficult to train: they either became unstable, entered a chaotic regime, or were unable to produce the desired movements altogether. Thus, we excluded ***h***(*t*) for all these networks, as well as for the 10 other ISNs that we constructed as part of this multi-network comparison.

##### ⋆2.1.5 Similarity transformation of network solutions

A trained (linear) network’s solution is completely determined by the triplet 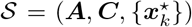, where 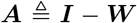 is the state transition matrix, ***C*** is the linear readout, and 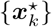 for *k* = 1,⋯, 8 is the set of movement-specific initial conditions. Using an invertible transformation 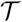, we can construct a new solution given by the triplet 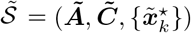, where

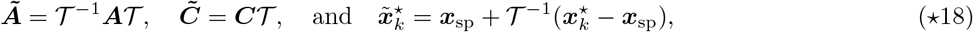

and 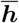 changed to 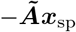 to ensure the spontaneous state remains the same. To see that 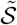 is also a solution to the task, we note that the output torques 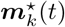 at any time *t* are equal to

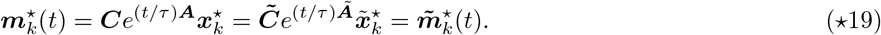

In Figure 6, we build the surrogate networks by applying the similarity transformation 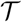 that (block-)diagonalizes ***A***, resulting in ***Ã*** of the form:

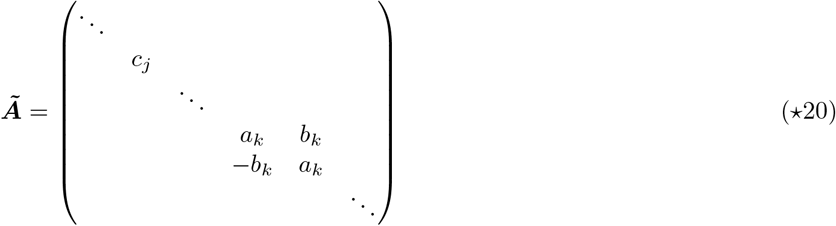

where {*c_k_*} and {*a_k_* ± *ib_k_*} are the sets of real and complex-conjugate eigenvalues of ***A***. The resulting ***Ã*** is a “normal” matrix (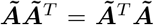; Trefethen and Embree, 2005), which—contrary to the original matrix ***A***—is unable to transiently amplify the set of movement-specific initial states 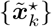 during movement (Figure 6D). In order to be able to meaningfully compare ║***C***║_F_ and 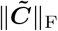 in Figure 6F, we scale 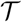 so as to preserve the average magnitude of the initial states (relative to the spontaneous state):

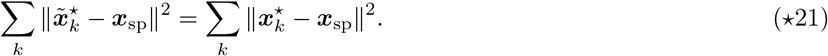

We can always achieve this because 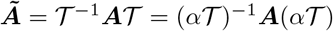 for any *α* ≠ 0.

#### ⋆2.2 Formalization of anticipatory motor control

We formalise the notion of anticipatory control by asking: given an intended movement (indexed by *k*), and the current (preparatory) state ***x***(*t*) of the network, how accurate would the movement be if it were to begin *now*? We measure this prospective motor error as the total squared difference 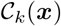 between the timecourses of the target torques ***m***⋆ and those that the network would generate (Equations ⋆8 and ⋆10) *if left uncontrolled* from time *t* onwards, starting from initial condition ***x***(*t*):

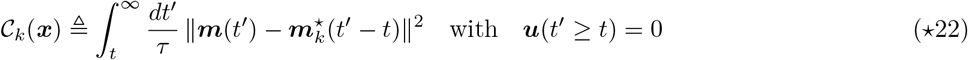

(we will often drop the explicit reference to the movement index *k* to remove clutter, as we did in the main text). Thus, any preparatory state ***x*** is associated with a prospective motor error 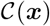.

The prospective error 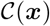 changes dynamically during movement preparation, as ***x***(*t*) evolves under the action of control inputs. The aim of the control inputs is to rapidly decrease this prospective error, until it drops below an acceptably small threshold, or until movement initiation is forced. We formalize this as the minimization of the following control cost:

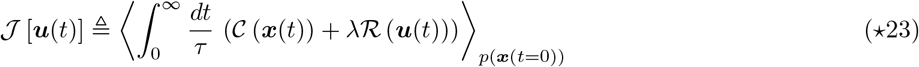

where 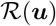 is a regularizer (see below) and the average 〈·〉 is over some distribution of states we expect the network to be found in at the time the controlled preparatory phase begins (we leave this unspecified for now as it turns out not to influence the optimal control strategy – see below). Thus, we want control inputs to rapidly steer the cortical network into states of low 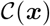 from which the movement can be readily executed, but these inputs should not be too large. The infinite-horizon summation expresses uncertainty about how long movement preparation will last, and indeed encourages the network to be “ready” as soon as possible.

Mathematically, 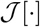 is a functional of the spatio-temporal pattern of control input ***u***(*t*) — indeed, ***x***(*t*) depends on ***u***(*t*) through Equation ⋆8. The regularizer 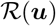, or “control effort”, encourages small control inputs and will be specified later. Without regularization, the problem is ill-posed, as arbitrarily large control inputs could be used to instantaneously force the network into the right preparatory state in theory, leading to physically infeasible control solutions in practice. Also note that Equation ⋆23 is an “infinite-horizon” cost, i.e. the integral runs from the beginning of movement preparation when control inputs kick in, until infinity. This does *not* mean, however, that the preparation phase must be infinitely long. In fact, good control inputs should (and will!) bring the integrand close to zero very fast, such that the movement is ready to begin after only a short preparatory phase (see e.g. Figure 4A in the main text).

In order to derive the optimal control law, we further assume that the dynamics of the network remain approximately linear during both movement preparation and execution. This is a good approximation provided only few neurons become silent in either phase (the saturation at zero firing rate is the only source of nonlinearity in our model, c.f. *ϕ*(·) in Equation ⋆8; Figure S5). In this case, the prospective motor error 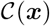 of Equation ⋆22 affords a simpler, interpretable form, which we derive now. In the linear regime, Equation ⋆8 becomes

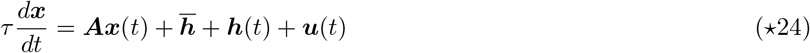

with an effective state transition matrix 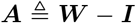. The network output at time *t*′, starting from state ***x*** at time *t*′ = 0 and with no control input thereafter, has an analytical form given by

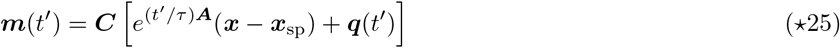

and similarly for ***m***⋆(*t*′) with ***x*** replaced by ***x***⋆. The final term ***q***(*t*′) is a contribution from the external input: it does not depend on the initial condition, and is therefore the same in both cases. Thus, the prospective motor error (Equation ⋆22) attached to a given preparatory state ***x*** is

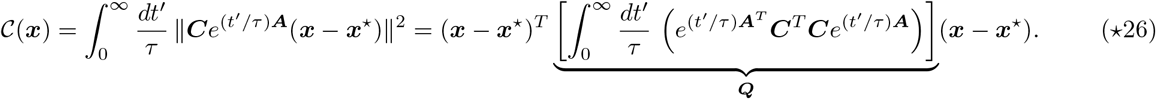

The matrix integral on the r.h.s. of Equation ⋆26 is known as the “observability Gramian” ***Q*** of the pair (***A, C***) (Skogestad and Postlethwaite, 2007; Kao and Hennequin, 2019). It is found algebraically as the solution to the Lyapunov equation^1^

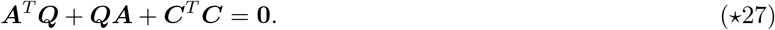

Thus, under linearity assumptions, the prospective motor error is a quadratic function of the difference between the momentary preparatory state ***x*** and an optimal initial state ***x***⋆ known to elicit the right muscle outputs in open loop. The Gramian ***Q***, a symmetric, positive-definite matrix, determines how preparatory deviations away from ***x***⋆ give rise to subsequent motor errors. Deviations along the few eigenmodes of ***Q*** associated with large eigenvalues will lead to large errors in muscle outputs. The optimal control input ***u***(*t*) will need to work hard to minimize this type of ‘prospectively potent’ deviations – luckily, there are few (see Figure 3D in the main text, where the top 20 eigenvalues of ***Q*** are shown; see also Supplementary Math Note 1 for further dissection). In contrast, errors occurring along eigenmodes of ***Q*** with small eigenvalues – the vast majority – have almost no motor consequences (‘prospectively null’). This large bottom subspace of ***Q*** provides a safe buffer in which preparatory activity is allowed to fluctuate without sacrificing control quality. It comprises both the “readout-null” and “dynamic-null” directions described in the main text (Figure 3). Geometrically, we can therefore think of the optimal preparatory subspace as a highdimensional ellipsoid centered on ***x***⋆, and whose small and (potentially infinitely) large axes are given by the top and bottom eigenvectors of ***Q***, respectively (small axes, steep directions, large eigenvalues; long axes, flat directions, small eigenvalues).

Finally, we note that the optimal control input ***u***(*t*) must keep the infinite-horizon integral in Equation ⋆23 finite. This is achieved if ***x***(*t*) reaches a fixed point equal to ***x***⋆, which is in turn achieved if the control input eventually settles in a steady state equal to

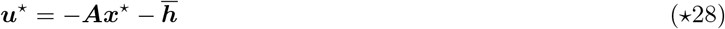

Thus, defining 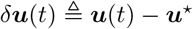 and 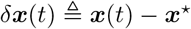, a relevant regularizer for our control problem is

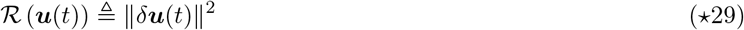

and our control cost functional becomes

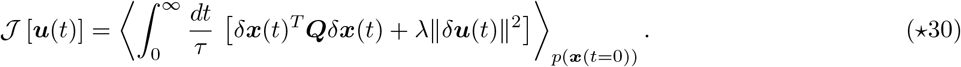

In our simulations, we perform a simple scalar normalization of ***Q*** so that Tr (***Q***) = *N*. This makes the first term of the cost more easily comparable to the energy penalty λ║*δ**u***║^2^, which also scales with *N*. In the next section, we show that the quadratic formulation of 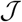 in Equation ⋆30 leads to analytically tractable optimization. In the rest of the Star Methods, as in the main text, we continue to assume linear network dynamics in order to derive optimal control laws but implement these solutions in the fully nonlinear circuit.

##### ⋆2.2.1 Classical LQR solution

When no specific constraints on ***u***(*t*) are imposed, the minimization of Equation ⋆30 is given by the celebrated linear quadratic regulator (LQR). Specifically, the optimal control input ***u***_opt_(*t*) = ***u***⋆ + *δ**u***_opt_(*t*) takes the form of instantaneous linear state feedback (Figure S3A):

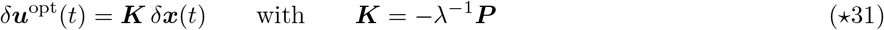

where ***P*** is a symmetric, positive-definite matrix, obtained as the solution to the following Riccati equation:

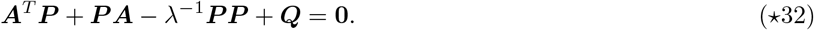

We will recover this optimal feedback law later (‘Feedback based on excitatory neurons only’) as part of a more general mathematical derivation; for now, we refer to standard texts, e.g. Skogestad and Postlethwaite, 2007.

Thus, to achieve optimal anticipatory control of fast movements, the best strategy for the preparatory phase is to feed back into the circuit a linearly weighted version of the momentary error signal *δ**x***(*t*). The optimal feedback matrix ***K*** turns out to not depend on the choice of distribution *p*(***x***(*t* = 0)). For a linear model, this also implies that ***K*** does not depend on the specific movement to be performed, i.e. on the specific state ***x***⋆ to be approached during preparation. Only the steady-state control input ***u***⋆ in Equation ⋆28 is movement-specific.

The total control cost is given by 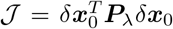, where *δ**x***_0_ is the deviation of the network state from ***x***⋆ at the onset of movement preparation (i.e., the state of the network at preparation onset is ***x***⋆ + *δ**x***_0_). The corresponding total energy cost 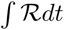 is given by 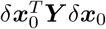 where ***Y*** is the solution to

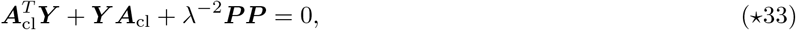

and

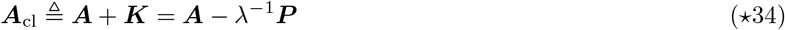

is the effective state matrix governing the dynamics of the closed control loop. The associated integrated prospective motor cost is then given by 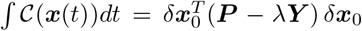. This is how we evaluate the total energy cost and integrated prospective motor cost in Figure 5H.

We emphasize that the LQR problem (and its solution in Equation ⋆31) assumes linear dynamics throughout the preparatory and movement epochs to compute (and optimize) the prospective motor cost 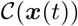. We verified that the quadratic form of Equation ⋆26 provides a good approximation to the actual 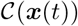 resulting from nonlinear dynamics during the movement epoch (Figure S5C). Moreover, all our simulations use the nonlinear activation function *ϕ*(·) in Equation ⋆8 unless indicated otherwise.

##### ⋆2.2.2 Naive feedforward solution

A straightforward solution exists for ensuring that, after enough preparation time, ***x***(*t*) converges exponentially to ***x***⋆ – thus *eventually* leading to the correct movement. This “naive” solution consists in setting ***u***(*t*) to the constant vector ***u***⋆ in Equation ⋆28 (thus *δ**u***(*t*) = 0 throughout preparation). Note that for the full nonlinear model, 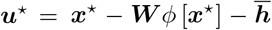. This constant input is provided during movement preparation and removed at the desired time of movement onset. This naive strategy does not rely on feedback, and so can be seen as a type of feedforward preparatory control. It also corresponds to the LQR solution in the limit of infinite energy penalty λ in Equation ⋆30, and indeed the solution in this limit yields ***K*** = 0, resulting in 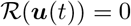.

#### ⋆2.3 Adapting optimal control to chaotic networks

In Figure 5D, we adapt optimal preparatory control to a chaotic network and find that LQR stabilizes its otherwise unstable dynamics during movement preparation. Thus far, we have assumed that ***A*** is Hurwitz-stable in deriving the optimal control law, which allows us to evaluate the integral in Equation ⋆26 analytically by solving Equation ⋆27. This integral diverges for unstable ***A***, and thus cannot be evaluated for a chaotic network. In this work, we simply set ***Q*** = ***I*** for the chaotic network, which likely overestimates the number of state-space directions that matter for preparatory control. More sophisticated methods could potentially be used for estimating the sensitivity of these nonlinear dynamics to errors in the initial condition. This would presumably lead to faster preparation for a set energy budget 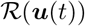, but we leave this for future work.

#### ⋆2.4 Preparing in the nullspace

In Figure S7, we extend the optimal control model to explicitly penalize non-zero output torques during movement preparation. Specifically, we augment the cost functional in Equation ⋆30 with a penalty on the integrated squared magnitude of ***m***(*t*), which can be written as a quadratic form in *δ**x*** much like the prospective motor cost in Equation ⋆26:

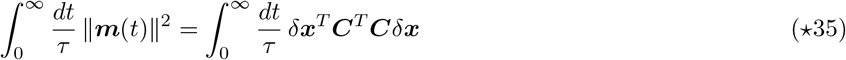

where ***m***(*t*) = ***C**δ**x*** because we have constrained the initial conditions 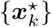 to be in the nullspace of ***C***. We thus write the combined cost functional as

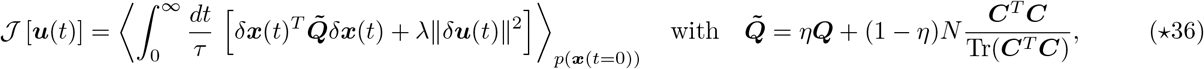

where *η* ∈ [0,1] is a scalar that weighs the relative importance of preparing fast and preparing while not moving (*η* = 0.2 in Figure S7).

#### ⋆2.5 Optimality under neural constraints

The linear quadratic regulator presented in ‘Classical LQR solution’ above brings the fundamental insight that control can (and in fact, should) be achieved via a feedback loop (Figure S3A). Such a loop could technically be embedded directly as a modification of the recurrent connectivity within M1, as all that matters for the control cost is the effective closed-loop state matrix ***A*** + ***K***. However, this would make it very difficult to switch the loop ON when movement preparation must begin, and OFF again when the movement is triggered. A more flexibly strategy would be to have the loop pass through another brain area, and gain-modulate this area (e.g. via inhibitory drive) to close or open the loop when appropriate.

A natural candidate structure for mediating such cortico-cortical feedback is the motor thalamus, which has been shown to be causally involved in movement preparation (Guo et al., 2017). Importantly, basic anatomy and physiology pose constraints on the type of connectivity and dynamics around the control loop, such that we will have to adapt the classical LQR theory to derive plausible circuit mechanisms. In particular, the thalamus is not innervated by the local inhibitory interneurons of M1, so feedback will have to computed based on the activity of (some of) the excitatory cells only, precluding full-state feedback. Moreover, the optimal LQR gain matrix ***K*** (given by Equations ⋆31 and ⋆32) contains both positive and negative elements with no structure; this violates Dale’s law, i.e. that neurons can be either excitatory or inhibitory but are never of a mixed type. Finally, the classical LQR solution prescribes *instantaneous* state feedback, whereas thalamic neurons will have to integrate their inputs on finite timescales, thereby introducing some “inertia”, or lag, in the feedback loop. In the rest of this section, we flesh out these biological constraints in more detail, and show that all of these limitations can be addressed mathematically, to eventually yield optimal control via a realistic thalamocortical feedback loop (see Figure S3B-D for a graphical overview).

##### ⋆2.5.1 Feedback based on excitatory neurons only

Here, we incorporate the key biological constraints that feedback from the cortex onto itself via the thalamus will have to originate from the excitatory cells only. Thus, instead of *δ**u***(*t*) = ***K**δ**x***(*t*), we look for a feedback matrix of the form (Figure S3B)

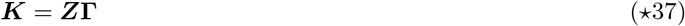

where 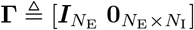 singles out the activity of the E neurons when computing the control input ***K**δ**x***, and ***Z*** is an *N* × *N*_E_ matrix of free parameters. To gain generality (which we will need later), we also assume that the control input enters the network through a matrix ***B***, i.e. the closed-loop state matrix (Equation ⋆34) becomes ***A***_cl_ = ***A*** + ***BZ*Γ**, and the energy penalty becomes λ║***B**δ**u***(*t*)║^2^. We now derive algebraic conditions of optimality for ***Z***, along with a gradient-based method to find the optimal ***Z*** that fulfills them.

First, we use the general result of Equation ⋆27 (see also Footnote 1) to rewrite the cost function 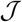 in Equation ⋆30 as:

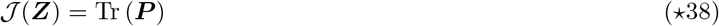

where ***P*** satisfies

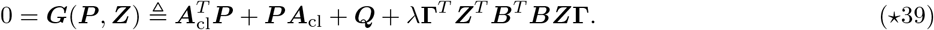

Note that 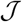 in Equation ⋆38 is now a function of the feedback matrix ***K***, and therefore of the parameter matrix ***Z***.

To minimize 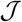 w.r.t. ***Z*** subject to the constraint in Equation ⋆39, we introduce the Lagrangian:

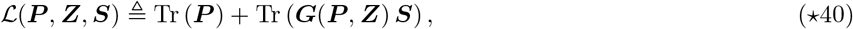

where ***S*** is a symmetric matrix of Lagrange multipliers (the matrix equality in Equation ⋆39 is symmetric, thus effectively providing *N*(*N* + 1)/2 constraints). After some matrix calculus, we obtain the following coupled optimality conditions:

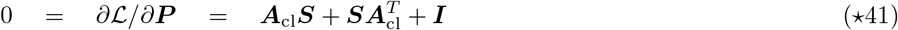

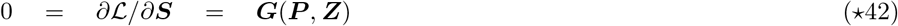

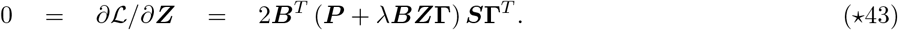

When the two Lyapunov equations Equations ⋆41 and ⋆42 are satisfied, the second term (Tr(***GS***)) in 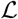 vanishes, such that 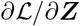 of Equation ⋆43 is in fact the gradient of Tr(***P***) w.r.t. ***Z*** subject to the algebraic constraint of Equation ⋆39. We use this gradient equation, together with the L-BFGS optimizer (Byrd et al., 1995) to find the optimal parameter matrix ***Z***. We then recover the optimal feedback gain matrix ***K*** according to Equation ⋆37. We start each optimization by setting 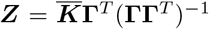, where 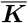 is the classical LQR solution to the same problem, such that ***Z*** is the least-square solution to 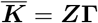.

##### ⋆2.5.2 Dale’s law

The previous subsection showed how to obtain a gain matrix ***K*** of size *N* × *N*_E_ that implements optimal, instantaneous cortico-cortical feedback originating from the excitatory cells. However, this optimal matrix typically has a mix of positive and negative elements that are not specifically structured. To implement the more realistic feedback architecture shown in Figure S3C, implicating the motor thalamus and M1 layer 4 (M1-L4), we seek a decomposition of the form

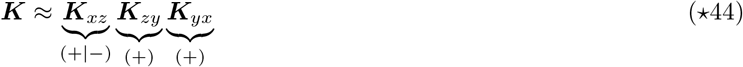

where ***K***_*yx*_ (M1 to thalamus) is an *N*_E_ × *N* matrix of non-negative elements, ***K***_*zy*_ (thalamus to M1-L4) is an *M* × *N*_E_ matrix of non-negative elements, and ***K***_*xz*_ (M1-L4 to the recurrent M1 network) is an *N* × *M* matrix composed of *M*_E_ non-negative columns and *M*_I_ non-positive columns (thus *M* = *M*_E_ + *M*_I_). Such a sign-structured decomposition will allow optimal control to be performed through the more realistic feedback architecture shown in Figure S3C, with corresponding dynamics of the form:

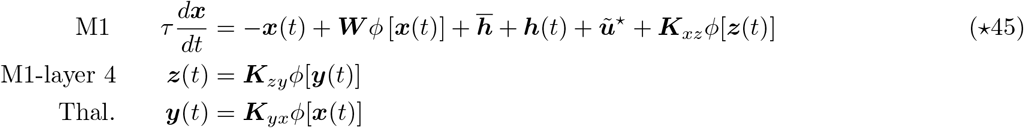

where

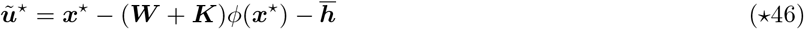

is a condition-dependent steady input given to the network during movement preparation so as to achieve the desired fixed point ***x***⋆.

To achieve this decomposition, we note that without loss of generality we can choose 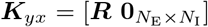 – where ***R*** is a random, element-wise positive *N*_E_ × *N*_E_ matrix – and apply the algorithm developed above (‘Feedback based on excitatory neurons only’) now with **Γ** = ***K***_*yx*_. This will return an optimal *N* × *N*_E_ matrix ***Z*** describing feedback from thalamus back to M1, which – as long as ***R*** is invertible – will achieve the same minimum cost as if ***R*** had been set to ***I***_*N*_E__. Here, we simply draw each element of ***R*** from Bernoulli(*p*), i.e. random sparse projections (the magnitude of ***R*** does not matter at this stage, as only the product ***Z*Γ** does; ***R*** will be renormalized later below). We now need to decompose this optimal feedback matrix as ***Z*** = ***K***_*xz*_ ***K**_zy_*, with the same sign constraints as in Equation ⋆44. We approach this via optimization, by minimizing the squared error implied by the decomposition, plus an 2-norm regularizer:

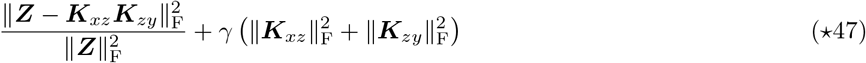

We parameterize each element of ***K**_xz_* and ***K**_zy_* as ±*z*^2^, where *z* is a free parameter to be optimized, and the ± sign enforces the sign structure written in Equation ⋆44. Minimization is achieved using L-BFGS and typically converges in a few tens of iterations. We note that the product ***K**_xz_**K**_zy_**K**_yx_* is invariant to any set of rescalings of the individual matrices as long as they cancel out to 1. Thus, after optimization, we re-balance the three matrices such that they have identical Frobenius norms. This is mathematically optional, but ensures that firing rates in M1, thalamus and M1-L4 have approximately the same dynamic range.

Importantly, we find that as long as the number of M1-L4 neurons (*M*) is chosen sufficiently large, the decomposition of ***Z*** that we obtain is almost exact, which implies that the dynamics of Equation ⋆45 still achieves optimal anticipatory control of movement under the architectural constraint of Equation ⋆37.

##### ⋆2.5.3 Taking into account integration dynamics in thalamus and M1-layer 4

The optimal control solution that we arrived at in Equation ⋆45 still relies on instantaneous feedback from cortex back onto itself. However, neurons in the thalamus and in M1’s input layer have their own integration dynamics – this will introduce lag around the loop, which must be taken into account when designing the optimal feedback. We therefore include these dynamics:

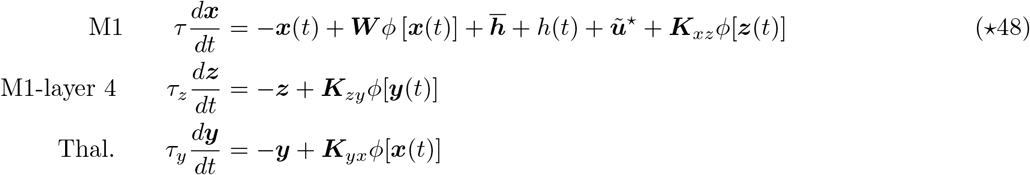

where the steady input 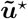 is again given by Equation ⋆46, and {*τ_y_, τ_z_*} are the single-neuron time constants in the thalamus and the cortical input layer. We then seek the optimal connectivity matrices {***K**_xz_, **K**_zy_, **K**_yx_*} to fulfill the same optimal-control principles as before, namely the minimization of the cost functional in Equation ⋆30. In order to do that, we note that the dynamics of ***x*** (M1 activity) in the linear regime do not change if the system of differential equations in Equation ⋆48 is simplified as

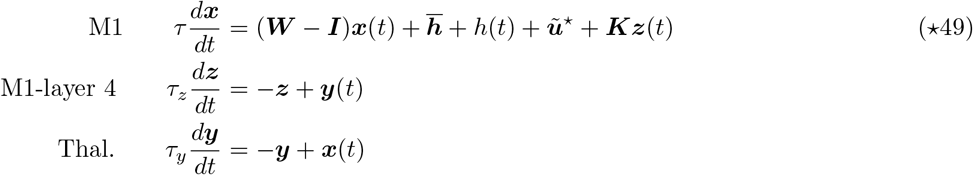

where ***K*** = ***K**_xz_ **K**_zy_ **K**_yx_* summarizes the three connectivity matrices around the loop into one effective feedback gain matrix. This formulation allows us to combine the steps developed in the previous two subsections to find the optimal connectivity matrices.

Specifically, we apply the algorithm developed in ‘Feedback based on excitatory neurons only’ to an augmented system with state matrix

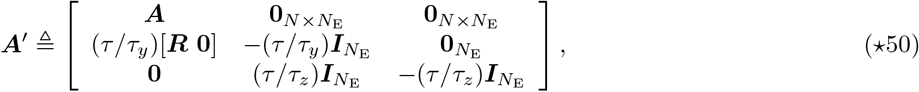

input matrix

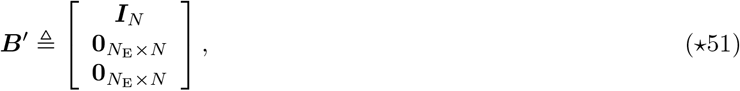

quadratic cost weighting matrix

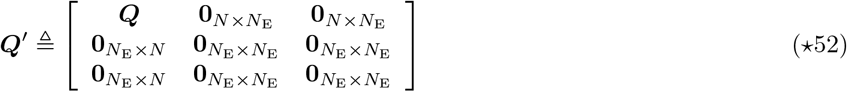

and feedback input parameterized as

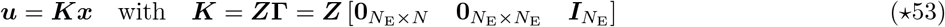

In Equation ⋆50, the matrix ***R*** is again a random matrix of sparse positive connections from M1 to thalamus.

The optimal ***Z*** (c.f. Equation ⋆43) corresponds to the product ***K**_xz_**K**_zy_*, which we can further decompose under sign constraints to recover the individual connectivity matrices ***K**_xz_* and ***K**_zy_*.

##### ⋆2.5.4 Disinhibitory action of the basal ganglia

We model the disinhibitory action of the basal ganglia (BG) on thalamic neurons as an on-off switch: to trigger movement, BG become active (BG neurons not explicitly modelled here) and the thalamic neurons are silenced instantly (i.e. ***y*** is set to **0**). When this happens, thalamic inputs to M1-L4 vanish and M1-L4 neural activity decays to zero on a time-scale *τ_z_* (see Equation ⋆50). As the activity of L4 neurons decays, these neurons continue to exert an influence on M1 activity through the connectivity matrix ***K**_zx_*. This lead to changes in movement-related M1 dynamics, resulting in small movement errors, which we correct post-hoc by re-optimizing the desired initial state ***x***⋆ for each movement. From these new desired states, network dynamics evolves—with the additional inputs from M1-L4 neurons after movement onset—to produce accurate hand trajectories. Crucially, unlike what we described in ‘Target hand trajectories and initial setup’, we do not re-optimize the readout matrix ***C*** here. This is because the observability Gramian ***Q*** and thus the closed-loop controller ***K*** depend on ***C***: changing the readout matrix ***C*** at this stage would cause the ***K*** we found to no longer be optimal with respect to 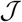. However, because the closed-loop solution does not depend on the desired fixed points ***x***⋆, we can re-optimize ***x***⋆ and still be guaranteed that the ***K*** that we found remains optimal.

##### ⋆2.5.5 Modelling the effect of photoinhibition

To model photoinhibition in our full circuit (whose dynamics are described by Equation ⋆45), we simply add a constant positive input *h*_ph_ to a subset of cortical inhibitory neurons chosen randomly, for a duration *T*_ph_ = 400 ms (see parameters in Table ⋆1). This results in an overall decrease in population activity across both excitatory and inhibitory neurons, consistent with the well-known paradoxical effects of adding positive inputs to I cells in inhibition-stabilized networks (Tsodyks et al., 1997; Ozeki et al., 2009; Sanzeni et al., 2019).

In Figure 8D, we consider how, after photoinhibition, the activity of the full circuit recovers in three subspaces: the coding subspace (CS), the persistent subspace (PS), and the remaining subspace (RS). To define the CS, we focus on *unperturbed* neural activity in the last 400 ms of movement preparation. For each movement condition, we compute the time-averaged population activity vector in this time window and combine them into a data matrix 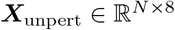. We perform principal component analysis (PCA) on **X**_unpert_ and extract an orthonormal basis ***U***_CS_, which captures 90% of the variance in ***X***_unpert_. The CS is defined as the subspace spanned by the columns of ***U***_CS_.

The PS and the RS are defined specifically for each perturbation experiment. In each experiment, we simulate the perturbed dynamics of the network for each movement condition, each with a different random subset of inhibitory neurons being ‘photostimulated’. To compute the PS, we consider the same time window that we do for the CS and calculate the *perturbed*, time-averaged population activity vector for each movement. We again collate them into a matrix 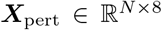 and perform PCA on the deviation induced by the perturbations in the given experiment, **Δ** = ***X***_pert_ – ***X***_unpert_. We extract an orthonormal basis ***U***_PS_, which captures 90% of the variance in **Δ**. The PS is defined as the subspace spanned by the columns of ***U***_PS_. Similarly, to compute the RS, we construct the data matrix ***Y*** = [***X***_unpert_ ***X***_pert_] in each perturbation experiment and orthogonalize it with respect to both ***U***_CS_ and ***U***_PS_. We perform PCA on the resulting data matrix and again extract an orthonormal basis ***U***_RS_ that captures 90% of the variance, which defines the RS. We find that the three subspaces combine to capture 98% of the total variance in ***Y***, averaged over 300 independent perturbation experiments.

To examine how activity trajectories recover in the three subspaces, we project the perturbed and unperturbed activity onto ***U***_CS_, ***U***_PS_, and ***U***_RS_ and calculate the magnitude of the deviation in each subspace. This is averaged over all movements and 300 independent perturbation experiments, which is what is shown in Figure 8D.

### ⋆3 Quantification and statistical analysis

#### ⋆3.1 Network measures

##### Participation ratio

To estimate the dimensionality of subspaces, we use the “participation ratio” (Gao et al., 2017), calculated based on the eigenvalue spectrum *σ*_1_, *σ*_2_,⋯, *σ_N_* of the relevant symmetric, positive-definite matrix as

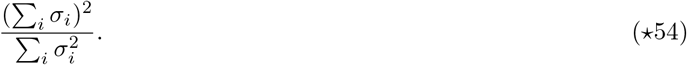

##### Nonnormality

In Figure 5E, we define the nonnormality of a network with synaptic connectivity matrix ***W*** as

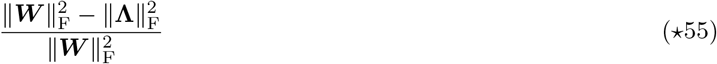

as proposed by Murphy and Miller (2009), where **Λ** is a diagonal matrix containing the eigenvalues of ***W***.

##### 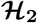 norm

In Figure 5F, we define the 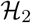 norm of a network as 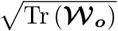, where the full-state observability Gramian 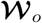 satisfies the Lyapunov equation 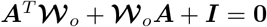 and ***A*** = −***I*** + ***W***.

##### Prospective motor potency

The prospective motor potency of a state space direction ***d*** (with ║***d***║ = 1) is ***d***^*T*^ ***Qd***, where ***Q*** satisfies Equation ⋆27. This quantifies the prospective motor error induced by a deviation ***x***⋆ + ***d*** of the final preparatory state away from ***x***⋆. In Figure 3D, we plot the top 20 eigenvalues of ***Q***, i.e. the motor potency of the 20 most potent directions (the corresponding eigenvectors of ***Q***). The prospective motor potency of a *K*-dimensional subspace 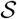, spanned by the unit vectors ***d***_1_,⋯, ***d**_K_*, is given by

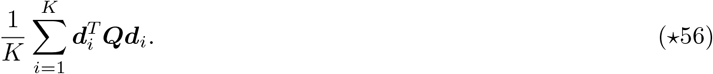

##### Amplification factor

In Figure 6D, for each movement *k* we define the amplification factor in the model as 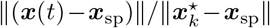, where ***x***(*t*) is the movement-epoch activity evolving according to Equation ⋆8 with ***h***(*t*) = 0, ***u***(*t*) = 0, and 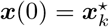. Description of how we define the amplification factor for the monkeys can be found below.

#### ⋆3.2 Neural data analysis

We analyzed neural recordings of two monkeys (J and N) performing a delayed reaching task (data courtesy of Mark Churchland, Matt Kaufman and Krishna Shenoy). Both the task and dataset have been described in detail previously (Churchland et al., 2010). Briefly, the two monkeys performed center-out reaches on a fronto-parallel screen. At the beginning of each trial, they fixated on the centre of the screen for some time, after which a target appeared on the screen. A variable delay period (0–1000 ms) ensued, followed by a go cue instructing the monkeys to reach towards the target. In this paper, we analyzed nine movement conditions, corresponding to the straight reaches that were most similar to the ones we modelled (Figure 2B). Moreover, we restricted our analysis to the trials with delay periods longer than 400 ms.

Recordings were made in the dorsal premotor and primary motor areas. We preprocessed the spike trains of 123 neurons for monkey J and 221 neurons for monkey N, following the same procedure outlined in Churchland et al. (2012). Briefly, we computed the average firing rates for each movement condition, further smoothed using a 20 ms Gaussian filter. Firing rates were computed separately for the delay and movement periods, time-locked to target and movement onset respectively; this is necessary because of variable delay periods and reaction times.

There is some subtletly in our definition of the time of movement onset in the model, when we compare its activity to monkey data. In the model, movement begins at the same time neural activity begins to undergo rapid changes, i.e. as soon as control inputs are removed. However, in the two monkeys, such rapid changes in neural activity occur roughly 100 ms before movement begins. We attribute such delays in movement to delays in downstream motor processes not considered in our model. Therefore, to align the temporal profile of neural activity in the model and the data, we define the time of “movement onset” in the model to be 100 ms after the control inputs are removed. We perform such temporal alignment in all the data and model comparisons shown in Figure 2, Figure 4, and Figure 6.

##### ⋆3.2.1 Overlap between preparatory end-states

We calculated the Pearson correlation across neurons between the preparatory end-states in both model and monkey data for all reaches. Preparatory end-states are defined as the activity states reached at the end of movement preparation (monkey activity aligned to the go cue). In both model and monkey data, preparatory end-states are similar (Figure 2B, bottom) for reaches with similar hand trajectories (hand trajectories Figure 2B, top), but negatively correlated for more distant movements.

##### ⋆3.2.2 Amplification factor

To compute the amplification factor for the two monkeys in Figure 6D, we consider firing rates in a 400 ms window starting 250 ms prior to movement onset. We remove the mean across conditions and compute for each reach condition, the manitude of the population activity vector, normalized by its magnitude at the start of this epoch. We then average this across all reach conditions. The amplification factor quantifies the expansion of the population activity across neurons during the movement epoch.

##### ⋆3.2.3 jPCA

We used the method described in Churchland et al. (2012) to identify state-space directions in which activity trajectories rotate most strongly. Briefly, we used numerical optimization to fit a skew-symmetric linear dynamical system of the form ***ẋ*** = ***Sx*** that best captures the population activity in a 400 ms window starting 220 ms before movement onset. We projected population activity in this window onto a plane spanned by the top two eigenvectors of ***S*** (Figure 2B).

##### ⋆3.2.4 Alignment index

To calculate the alignment index, we closely followed the methods described in Elsayed et al. (2016). The alignment index is defined as the (normalized) percentage of across-condition variance during movement captured by the top *K* principal components (PCs) of the preparatory activity:

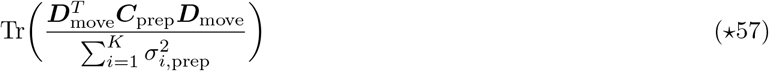

where the *K* columns of ***D***_move_ are the top *K* principal components of move. activity (“move-PCs”), ***C***_prep_ is the covariance matrix of prep. activity, and 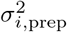 is the prep. activity variance captured by the *i*^th^ prep-PC. We choose *K* such that *K* prep-PCs captures 80% of the variance in prep. activity. Here, we define prep. activity as the delay-period activity during a 300 ms window starting 150 ms after target onset; the activity is calculated time-locked to target onset. Similarly, move. activity is defined as activity during a 300 ms window starting 50 ms prior to movement onset; the activity is calculated using firing rates time-locked to movement onset.

Methods for calculating the control of the alignment index are described in detail in the Supplementary Material of Elsayed et al. (2016) and are not reproduced here. For the model, the alignment index is calculated in the same way as for the neural data.

##### ⋆3.2.5 Canonical-correlation analysis

To compare model and monkey activity, we performed canonical-correlation analysis (CCA) on activity in a time window starting 400 ms before and ending 400 ms after movement onset (see discussion above for nuance in defining the time of movement onset in the model). To avoid overfitting to noise in CCA (Sussillo et al., 2015; Raghu et al., 2017), we first reduced the dimensionality of the two data sets, by projecting activity onto the top 21 (monkey J), 21 (monkey N), and 14 (model) principal components; the number of principal components are chosen to capture 95% of the across-condition activity variance in the two datasets. We then calculated the canonical correlations between the two reduced data sets, using the numerically stable algorithm described in Press (2011). We found that monkey and model activity are similar across time and reaches, with a high average canonical correlation (Figure 2C-E). We obtained similar results when we varied the number of principal components kept in the two data sets, which in turn varied the number of canonical variables.

### ⋆4 Additional resources

None.

### ⋆5 Key resources table

★5 Key resources table
| REAGENT OR RESOURCE | SOURCE | IDENTIFIER |
| --- | --- | --- |
| Software and Algorithms |  |  |
| OCaml 4.10 | Open source | http://www.ocaml.org |
| Owl | Open source | http://ocaml.xyz |
| Python 3 (monkey data preprocessing) | Open source | http://python.org |
| Gnuplot 5.2 | Open source | http://gnuplot.info |

The values of all the parameters used in this study are listed in the table below.

## Footnotes

1 This result is central to the theory of linear quadratic control, where cost functions are often of the form of integrated squared functions of the state, output, or input, under linear dynamics. It allows one to manipulate these integrals algebraically, and compute them numerically by solving a linear matrix equation (e.g. Bartels and Stewart, 1972). Indeed we will use this result again several times below in different contexts.

