## Supplementary Material for "Optimal anticipatory control as a theory of motor preparation: a thalamo-cortical circuit model"

### Inventory

**1 Supplementary Math Note: Control geometry of the ISN (related to Figures 3 and 4) 2**

**2 Supplementary Math Note: Universal performance of naive feedforward control (related to Figure 5) 3**

**3 Supplementary Math Note: Alignment index under naive feedforward control (related to Figure 6) 3**

**Supplementary Figures 5**

S1 Arm kinematics and target reaches (related to Figure 2) . . . . . . . . . . . . . . . 5
S2 Comparison of movement-generating network model with monkey M1 dynamics during movement (related to Figure 2) . . . . . . . . . . . . . . . . . . . . . . . . . 6
S3 Biologically plausible implementation of optimal anticipatory motor control (related to Figure 7) . . . . . . . . . . . . . . . . . . . . . . . . . . . . . . . . . . . 7
S4 Control geometry of the ISN (related to Figures 3 and 4) . . . . . . . . . . . . . . 7
S5 Role of the nonlinearity in preparatory control (related to Figure 4) . . . . . . . 8
S6 Comparison of movement-generating dynamics in two other classes of trained networks (full and low-rank) and in monkey data (related to Figure 5) . . . . . . 9
S7 Preparing whilst not moving (related to Figures 4 and 6) . . . . . . . . . . . . . 9

### 1 Supplementary Math Note: Control geometry of the ISN (related to Figures 3 and 4)

The optimal LQR strategy can be used to expose the challenges associated with controlling the inhibition-stabilized model of M1 that we use here. Indeed, network activity may be more easily controlled (or "steered") along some directions than along others, and having analytical access to the optimal control inputs (Equations ⋆31 and ⋆32) allows us to quantify this "control geometry". Specifically, we quantify control performance as $\mathcal{E} = \int_0^\infty (\delta \boldsymbol{x}^T \boldsymbol{Q} \delta \boldsymbol{x}) dt$, i.e. our original cost functional $\mathcal{J}$ in Equation ⋆23 without the input energy penalty. We can then ask: what is the smallest such cost $\mathcal{E}_{\text{min}}$ that can be achieved with a fixed input energy budget $\int_0^\infty \|\delta \boldsymbol{u}\|^2 dt$? We know that $\mathcal{E}_{\text{min}}$ is achieved by the LQR solution $\delta \boldsymbol{u} = \lambda^{-1} \boldsymbol{P}_\lambda$ (we use the $\cdot_\lambda$ subscript to make the dependence of $\boldsymbol{P}$ on $\lambda$ explicit). It can be shown that the input energy induced by this optimal feedback law is a decreasing function of $\lambda$. Thus, all we need to do is find the $\lambda$ that gives us the desired value of $\int_0^\infty \|\delta \boldsymbol{u}\|^2 dt$, and evaluate $\mathcal{E}_{\text{min}}$ for this particular $\lambda$. Importantly, the result will depend on the state of the cortical network at the beginning of the controlled preparatory phase, relative to the target $\boldsymbol{x}^\star$.

A simple derivation based on the LQR Ricatti equation shows that starting the control phase from some initial condition $\boldsymbol{x}^\star + \delta \boldsymbol{x}_0$ yields a total control cost equal to $\mathcal{J} = \delta \boldsymbol{x}_0^T \boldsymbol{P}_\lambda \delta \boldsymbol{x}_0$. Moreover, the corresponding energy cost is given by $\delta \boldsymbol{x}_0^T \boldsymbol{Y} \delta \boldsymbol{x}_0$ where $\boldsymbol{Y}$ is the solution to

$$\boldsymbol{A}_{\text{cl}}^T \boldsymbol{Y} + \boldsymbol{Y} \boldsymbol{A}_{\text{cl}} + \lambda^{-2} \boldsymbol{P}_\lambda \boldsymbol{P}_\lambda = 0, \tag{S1}$$

and

$$\boldsymbol{A}_{\text{cl}} \triangleq \boldsymbol{A} + \boldsymbol{K} = \boldsymbol{A} - \lambda^{-1} \boldsymbol{P}_\lambda \tag{S2}$$

is the effective state matrix governing the dynamics of the closed control loop. For a given $\delta \boldsymbol{x}_0$, we use a simple root-finding method (bisection with initial interval bracketting) to find the $\lambda$ that achieves the set, desired energy cost (our fixed "energy budget"). For this $\lambda$, we then calculate the associated control cost $\mathcal{E} = \delta \boldsymbol{x}_0^T (\boldsymbol{P} - \lambda \boldsymbol{Y}) \delta \boldsymbol{x}_0$. This is plotted in Figure S4, for initial deviations of $\boldsymbol{x}$ from $\boldsymbol{x}^\star$ chosen to be the top 20 eigenvectors of $\boldsymbol{Q}$, ranked by their respective eigenvalues $\nu_i$ (Equation ⋆26).

Figure S4 (right) shows that there is "no free lunch": preparatory deviations from $\boldsymbol{x}^\star$ that induce the worst motor errors (the top eigenvectors of $\boldsymbol{Q}$, with the largest eigenvalues $\nu_i$) are also those that are the most difficult to control, i.e. for which the minimal control cost $\mathcal{E}_{\text{min}}$ will be largest for a fixed input energy budget.

From the point of view of dynamical systems, this result is rather intuitive. The optimal initial conditions $\{\boldsymbol{x}_k^\star\}$ (found via optimization to achieve the required torques; Section ⋆2.1) are positioned in state space where the flow induced by the recurrent connectivity is strong – strong enough to elicit rich transients that can be decoded into torques patterns that grow transiently before decaying. To steer M1 towards (and maintain it at) these states, the input $\delta \boldsymbol{u}(t)$ (and the steady input $\boldsymbol{u}^\star$) must work against the strong local flow of the recurrent dynamics. This requires large inputs. From a physiological standpoint, this is also intuitive. The optimal initial states $\{\boldsymbol{x}_k^\star\}$ are shown to be states in which the E/I balance is momentarily broken (Hennequin et al., 2014). Much input energy must be spent to sustain an E/I imbalance in a network whose connectivity strives to maintain balance.

#### 2 Supplementary Math Note: Universal performance of naive feedforward control (related to Figure 5)

In this section, we show that for a linear system solving a motor task, the anticipatory control cost $\mathcal{J}$ under the naive feedforward strategy only depends on the movements that are generated as part of the task.

Consider a network characterized by a state matrix $\boldsymbol{A}$ and generating a given movement using readout weights $\boldsymbol{C}$ and an initial condition $\boldsymbol{x}^\star$ using the following movement-epoch dynamics:

$$\tau \frac{d\boldsymbol{x}}{dt} = \boldsymbol{A}\boldsymbol{x}(t) + \overline{\boldsymbol{h}} \tag{S3}$$

where $\overline{\boldsymbol{h}} = -\boldsymbol{A}\boldsymbol{x}_{\text{sp}}$ sets the spontaneous fixed point at $\boldsymbol{x}_{\text{sp}}$. These are the same dynamics as Equation ⋆8 but excluding $\boldsymbol{h}(t)$ as explained in Section ⋆2.1.4. When the feedforward strategy is used for preparatory control, a constant preparatory input $\boldsymbol{u}^\star$ (Equation ⋆28) is added to the r.h.s. of Equation S3 that instantaneously switches the network's fixed point from $\boldsymbol{x}_{\text{sp}}$ to $\boldsymbol{x}^\star$. Thus,

$$\boldsymbol{x}(t) = \boldsymbol{x}^\star + e^{\frac{t}{\tau}\boldsymbol{A}}(\boldsymbol{x}_{\text{sp}} - \boldsymbol{x}^\star). \tag{S4}$$

Now, let $\boldsymbol{x}(t, t')$ be the activity of the network at time $t+t'$, where $t$ marks the end of preparation and the beginning of the movement. At time $t$, the constant preparatory input $\boldsymbol{u}^\star$ is withdrawn, causing the fixed point of the dynamics to switch back to $\boldsymbol{x}_{\text{sp}}$. Therefore,

$$\boldsymbol{x}(t, t') = \boldsymbol{x}_{\text{sp}} + e^{\frac{t'}{\tau}\boldsymbol{A}}(e^{\frac{t}{\tau}\boldsymbol{A}} - I)(\boldsymbol{x}_{\text{sp}} - \boldsymbol{x}^\star) \tag{S5}$$

and the corresponding output torques as $\boldsymbol{C}\boldsymbol{x}(t, t')$. To compute the corresponding *error* in torques, $\boldsymbol{m}(t, t') - \boldsymbol{m}^\star(t')$, we note that the system produces the target movements with no error in the limit of infinite preparation time ($t \to \infty$). Thus, the momentary error at time $t + t'$ is given by

$$\begin{aligned} \boldsymbol{m}(t, t') - \boldsymbol{m}^\star(t') &= \boldsymbol{C}\left[\boldsymbol{x}(t, t') - \boldsymbol{x}(\infty, t')\right] && \text{(S6)} \\ &= \boldsymbol{C}e^{\frac{t+t'}{\tau}\boldsymbol{A}}(\boldsymbol{x}_{\text{sp}} - \boldsymbol{x}^\star) && \text{(S7)} \end{aligned}$$

where we have also used the fact that $\boldsymbol{C}\boldsymbol{x}_{\text{sp}} = 0$. Importantly, it is easy to show that Equation S7 is in fact equal to the target output torque $t + t'$ seconds after movement onset, i.e. $\boldsymbol{m}^\star(t + t') = \boldsymbol{C}\boldsymbol{x}(\infty, t + t')$. In summary, under the naive feedforward control strategy, motor errors following insufficiently long preparation only depend on the target movements, but not on the details of the (linear) system that achieves them. *A fortiori*, since the control cost $\mathcal{J}$ (Equation ⋆30) is defined based on output errors in torques, it must also be the same for any network that has been successfully trained to produce the target torques (in the limit of long preparation). We calculated this cost to be

$$\mathcal{J}_{\text{naive}} = \frac{1}{\tau^2} \iint_0^\infty dt\, dt' \, \|\boldsymbol{m}^\star(t + t')\|^2. \tag{S8}$$

#### 3 Supplementary Math Note: Alignment index under naive feedforward control (related to Figure 6)

Early preparatory activity under the naive feedforward strategy can be shown to be the negative image of movement-epoch activity, up to a constant offset. Consider a network solution to the reaching task characterized by linear dynamics with a state matrix $\boldsymbol{A}$ and a set of initial conditions $\{\boldsymbol{x}_k^\star\}$. During the preparatory epoch for movement $k$, the network activity evolves according to

$$\tau \frac{d\boldsymbol{x}}{dt} = \boldsymbol{A}\boldsymbol{x}(t) + \bar{\boldsymbol{h}} + \boldsymbol{u}_k^\star \tag{S9}$$

where $\boldsymbol{u}_k^\star$ sets the fixed point of the preparatory dynamics at $\boldsymbol{x}_k^\star$. Thus, preparatory activity $\boldsymbol{x}_k^{\mathrm{p}}(t)$ for a movement $k$ is given by:

$$\boldsymbol{x}_k^{\mathrm{p}}(t) = \boldsymbol{x}_k^\star + e^{\frac{t}{\tau}\boldsymbol{A}}(\boldsymbol{x}_{\mathrm{sp}} - \boldsymbol{x}_k^\star). \tag{S10}$$

Movement-epoch dynamics, on the other hand, obey:

$$\tau \frac{d\boldsymbol{x}}{dt} = \boldsymbol{A}\boldsymbol{x}(t) + \bar{\boldsymbol{h}} + \boldsymbol{h}(t) \tag{S11}$$

where $\bar{\boldsymbol{h}} = -\boldsymbol{A}\boldsymbol{x}_{\mathrm{sp}}$ sets the fixed point at $\boldsymbol{x}_{\mathrm{sp}}$ and $\boldsymbol{h}(t)$ is the condition-independent input ($\boldsymbol{h}(t) = \boldsymbol{0}$ for the 'full' and 'low-rank' networks). Assuming the network activity successfully reaches $\boldsymbol{x}_k^\star$ during the preparation epoch, movement activity $\boldsymbol{x}_k^{\mathrm{m}}(t)$ is given by

$$\boldsymbol{x}_k^{\mathrm{m}}(t) = \boldsymbol{x}_{\mathrm{sp}} + e^{\frac{t}{\tau}\boldsymbol{A}}(\boldsymbol{x}_k^\star - \boldsymbol{x}_{\mathrm{sp}}) + \boldsymbol{q}(t), \tag{S12}$$

where

$$\boldsymbol{q}(t) = \int_0^t e^{\frac{t-t'}{\tau}\boldsymbol{A}}\boldsymbol{h}(t')dt' \tag{S13}$$

is the contribution of the condition-independent external drive $\boldsymbol{h}(t)$. Comparing Equation S10 and Equation S12, we find that $\boldsymbol{x}_k^{\mathrm{p}}$ is the negative image of $\boldsymbol{x}_k^{\mathrm{m}}$, up to a constant offset, and condition-independent temporal variations:

$$\boldsymbol{x}_k^{\mathrm{p}}(t) = -\boldsymbol{x}_k^{\mathrm{m}}(t) + \boldsymbol{x}_k^\star + \boldsymbol{x}_{\mathrm{sp}} + \boldsymbol{q}(t). \tag{S14}$$

In computing the alignment index, Elsayed et al. (2016) first removed the mean across condition at every time point. We have done the same mean removal in our analysis, which eliminates the $\boldsymbol{x}_{\mathrm{sp}} + \boldsymbol{q}(t)$ terms in Equation S14, yielding

$$\hat{\boldsymbol{x}}_k^{\mathrm{p}}(t) = -\hat{\boldsymbol{x}}_k^{\mathrm{m}}(t) + \hat{\boldsymbol{x}}_k^\star \tag{S15}$$

where $\hat{\cdot}$ denotes deviation from the condition mean. This result explains the high alignment index of all network classes under the naive feedforward strategy (Figure 6C).

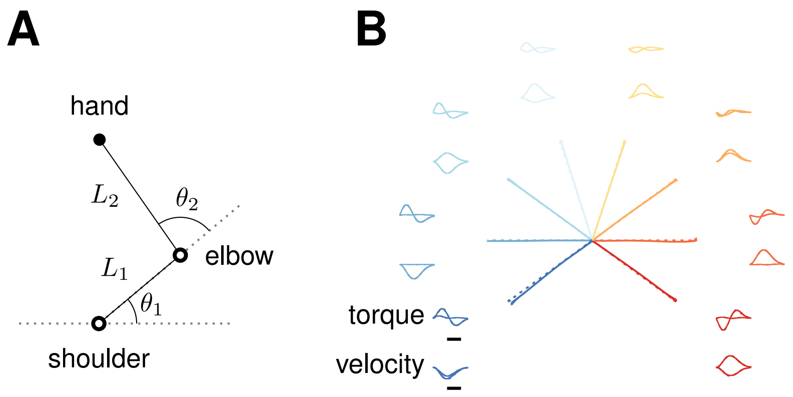


Figure S1: [Related to Figure 2] **(A)** Schematics of the arm model. **(B)** Reaches produced by the model, along with associated torques at the two joints, and x-y velocities of the hand (solid lines). Dashed lines denote target trajectories. Scale bar: 200 ms.

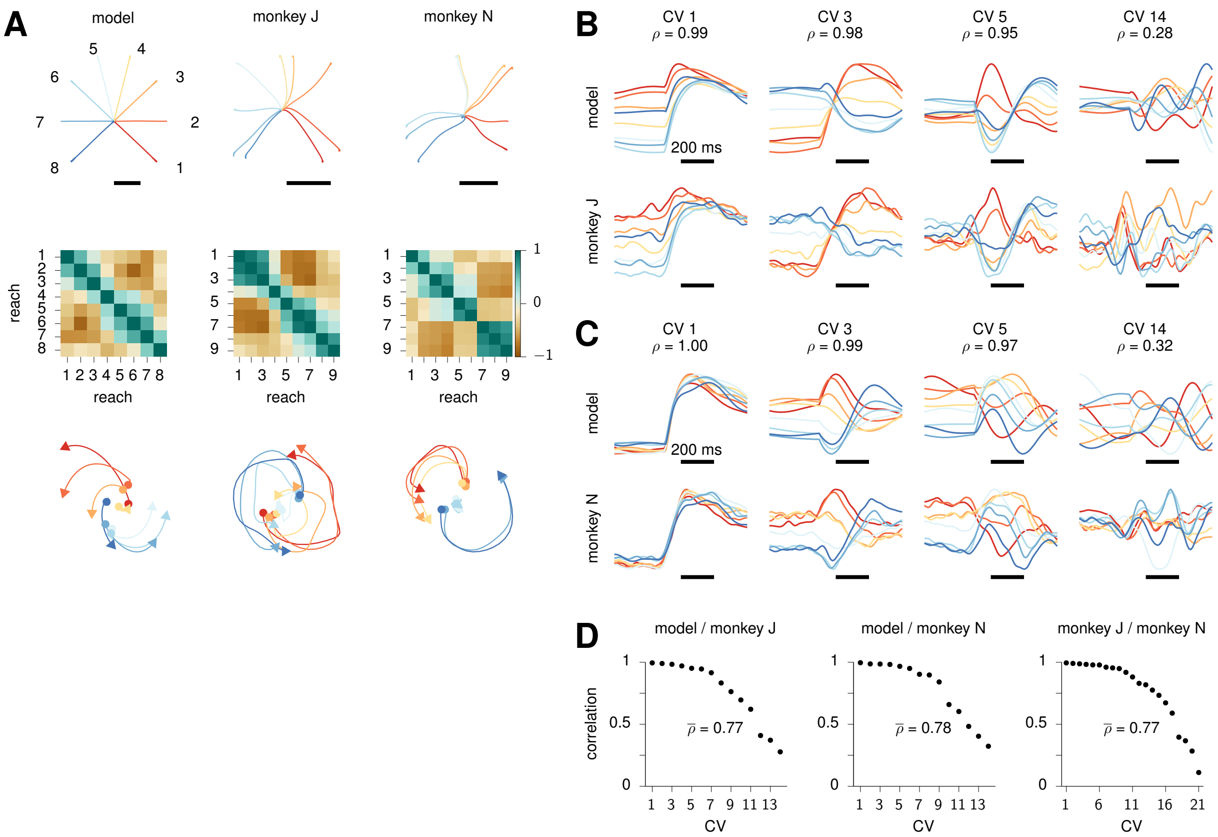


Figure S2: [Related to Figure 2] **Comparison of movement-generating network model with monkey M1 dynamics during movement.** **(A)** Top: model (left) and monkey (center and right) hand trajectories for each straight reach (color-coded). Black bars denote 10 cm. Monkey hand trajectories are averaged across trials with delays longer than 400 ms. Middle: overlap (Pearson correlation across neurons) between the preparatory end-states for model (left) and monkey activity (center and right). Reaches are numbered counter-clockwise as indicated near the model hand trajectories. Bottom: neural activity projected into the top jPC plane (see text). **(B)** Timecourse of the 1st, 3rd, 5th and 14th CCA projections (canonical variables) in the model (top) and monkey J (bottom), for each condition (color-coded as in B). Black scale bars indicate 200 ms from movement onset (note that "movement onset" in the model is re-defined to account for the latency between the go cue and actual movement onset in the monkey; see Section ⋆3.2). To equalize the number of movement conditions across model and monkeys, we dropped the 9th movement, which is kinematically redundant the 8th (c.f. B). **(C)** Same as C, for monkey N (5th movement excluded, redundant with the 6th). **(D)** Full spectrum of canonical correlations, with average labeled $\bar{\rho}$.

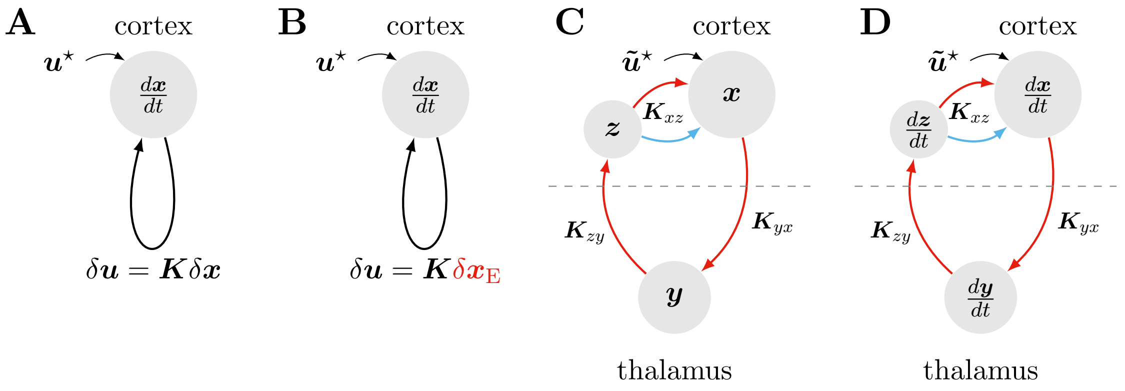


Figure S3: [Related to Figure 7] Four steps to arrive at a biologically plausible implementation of optimal anticipatory motor control. **(A)** The classical LQR solution prescribes instantaneous state feedback, with reentrant control inputs of the form of $\delta\boldsymbol{u}(t) = \boldsymbol{K}\delta\boldsymbol{x}(t)$ and a constant external input $\boldsymbol{u}^\star$. Section ⋆2.2.1 shows how to obtain the optimal feedback matrix $\boldsymbol{K}$ in this case. **(B)** It is possible to constrain feedback to be of the form $\delta\boldsymbol{u}(t) = \boldsymbol{K}\,[\boldsymbol{I}_{N_\mathrm{E}}\ \boldsymbol{0}_{N_i}]\,\delta\boldsymbol{x}(t) = \boldsymbol{K}\delta\boldsymbol{x}_\mathrm{E}(t)$ instead. Section ⋆2.5.1 shows how to obtain the optimal feedback matrix $\boldsymbol{K}$ in this case. **(C)** For flexibility, we propose that feedback be relayed by the motor thalamus, which is under the gating control of the basal ganglia. In Section ⋆2.5.2, we show that the optimal feedback gain $\boldsymbol{K}$ obtained in (B) can be decomposed into sign-constrained matrices implementing E connections from M1 to thalamus ($\boldsymbol{K}_{yx}$), from thalamus to M1-layer 4 ($\boldsymbol{K}_{zy}$), and Dale-structured E/I connections from layer 4 back into the main recurrent M1 circuit ($\boldsymbol{K}_{xz}$). **(D)** Finally, first-order dynamics can be introduced in our model thalamus and M1-layer 4 neurons. We show in Section ⋆2.5.3 how the lag introduced by such dynamics can be taken into account, to obtain a set of connections that achieve optimal anticipatory control of movement under these biological constraints.

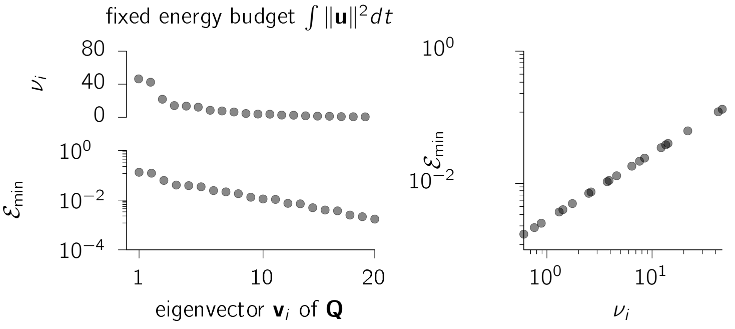


Figure S4: [Related to Figures 3 and 4] **Left:** eigenvalues $\nu_i$ (top) and minimum control cost $\mathcal{E}_\mathrm{min}$ achievable given a fixed energy budget (see text), for the top 20 eigenvectors of the observability Gramian $\boldsymbol{Q}$ defined in Equation ⋆26. Note that $\nu_i$ is also the motor error $\mathcal{C}$ incurred by a deviation of the preparatory state $\boldsymbol{x}^\star$ of unit length in the direction of the corresponding eigenvector $\boldsymbol{v}_i$, just prior to movement. **Right:** same $\nu_i$ and $\mathcal{E}_\mathrm{min}$ as shown on the left, plotted against each other.

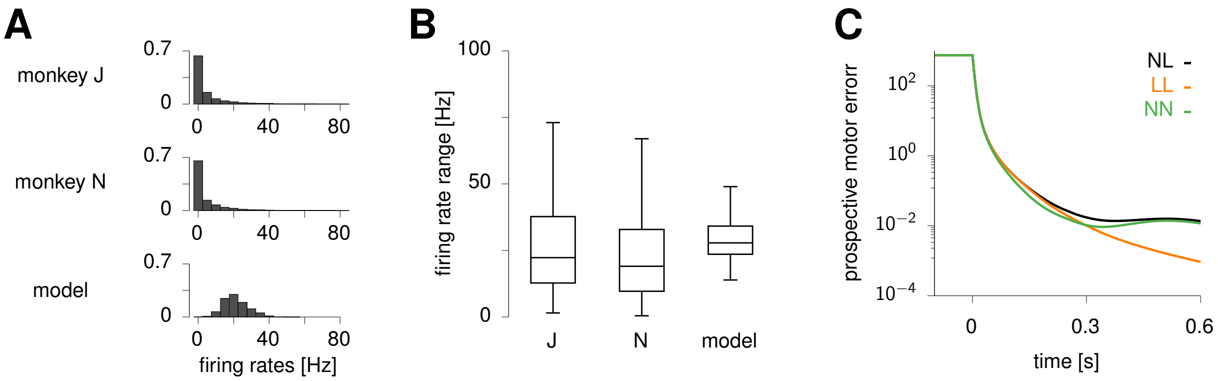


Figure S5: [Related to Figure 4] **Role of the nonlinearity in preparatory control.** **(A)** Distribution of firing rates across all neurons, times (including both preparation and movement epochs) and reach conditions. **(B)** Firing rate range (max − min) across time and reach conditions, computed for each neuron in monkey J, monkey N, and the model (min, 25th percentile, median, 75th percentile, max). **(C)** Prospective motor error during optimal preparation, under three different assumptions. NL: nonlinear network dynamics during preparation, but prospective error computed assuming linear dynamics during movement (c.f. Figure 4A of the main text). LL: same as NL, but with linear dynamics during preparation. NN: nonlinear dynamics during preparation, and prospective error computed using rollouts of the same nonlinear network dynamics during the movement-epoch. In all three cases, the optimal preparatory inputs were computed assuming a linear model as throughout the paper.

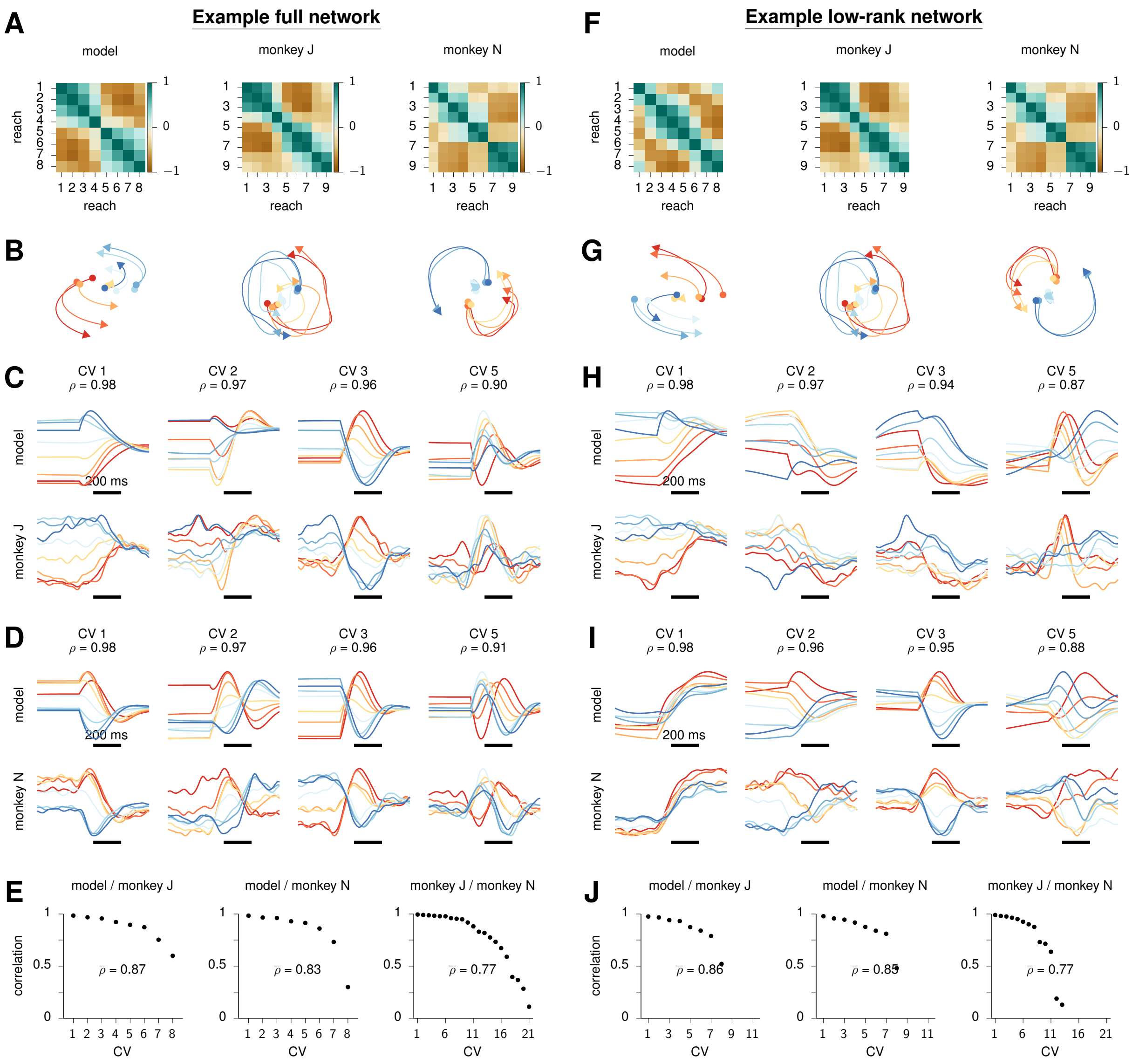


Figure S6: [Related to Figure 5] Movement-generating dynamics in an example 'full' network **(A-E)** and an example 'low-rank' network **(F-J)**, and comparison to monkey M1 dynamics. For details, please see the caption of Figure S2 where the same panels are shown for the ISN model.

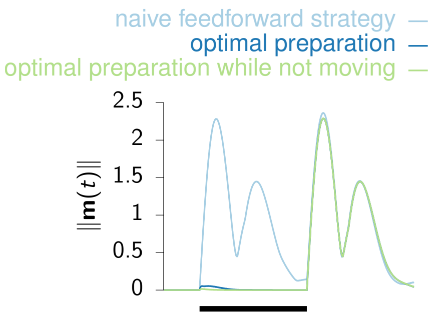


Figure S7: [Related to Figures 4 and 6] **Preparing whilst not moving.** Norm of the output joint torques, $\|\boldsymbol{m}(t)\|$, averaged over the 8 reaches, for the naive feedforward strategy (light blue), optimal preparatory control (dark blue), and an extended optimal preparatory control strategy that explicitly penalizes premature movement (green). The black bar marks the preparatory epoch.
